# Intermittent AP-1 activation in muscles contributes to exercise-induced health benefits

**DOI:** 10.64898/2026.01.05.696542

**Authors:** Abhinav Choubey, Udhaya Kumar S., Tingting Yang, Jennifer Jager, Sungguan Hong, Manashree Damle, Huiqi Yin, Ruxing Zhao, Chengchuang Song, Ege Sanem Oturmaz, Hui Tong, Jason Puglise, Elisabeth R. Barton, Bin Fang, Zhaoyong Hu, Lan Liao, Jianming Xu, Ye Lan, Zheng Sun

## Abstract

Regular physical exercise extends healthspan, yet the molecular mechanisms that translate intermittent contractile stress into lasting benefit remain incompletely understood. Using global nuclear run-on (GRO-seq) in mouse skeletal muscle after treadmill running, we profiled enhancer RNA (eRNA), a sensitive marker of enhancer activity. Activation protein-1 (AP-1), a family of pioneering factors for senescence, emerged as the top transcription factor with motif enrichment in exercise-activated enhancers. Our screen in the contracting C2C12 myotubes pinpointed cFos/JunD as the primary AP-1 factor responsible for contraction-induced transcriptional changes. Muscle-specific overexpression of A-Fos, a dominant-negative mutant of cFos, disrupted transcriptomic responses to exercise and attenuated exercise-mediated improvement in muscle functions. Interestingly, intermittent but not continuous overexpression of cFos/JunD in mouse muscles mimicked exercise-induced transcriptomic changes, increased mitochondrial volume density, enhanced muscle strength and fatigue resistance, and improved glucose tolerance. These results define a transcriptional regulatory signaling pathway linking exercise intermittency to beneficial adaptations and highlight the necessary recovery cycles in training. The paradoxical anti- and pro-aging roles of AP-1 offer insights into the timing and dynamics of stressors and stress responses in shaping senescence and healthspan.

## INTRODUCTION

Regular physical exercise can extend healthspan and counteract aging-associated conditions, such as muscle weakening, diabetes, and dyslipidemia^1,2^. Exercise brings multicentric adaptations that improve metabolic health through paracrine, autocrine, and endocrine mechanisms, acting at both cell-autonomous and non-autonomous levels. Skeletal muscle is the largest organ by mass and is particularly adaptive to exercise training^3,4^. Exercise counters age-related changes of skeletal muscles, including muscle mass, strength, endurance, mitochondrial morphology and function, insulin sensitivity, and intermediary metabolism.

Progress has been made in understanding the molecular mechanisms underlying exercise-induced benefits in muscles. For example, peroxisome proliferator-activated receptor γ coactivator 1α (PGC1α) gene expression is elevated in muscles after exercise^5–7^. However, muscle-specific overexpression of PGC1α promotes lipid accumulation, induces insulin resistance, reduces strength, and causes oxidative stress and inflammatory responses despite enhanced mitochondrial biogenesis ^8–11^. AMP-activated protein kinase (AMPK) is activated upon exercise-induced energy stress, helping cells conserve energy by inhibiting energy-consuming processes and promoting energy production.Its long-term effects are partly mediated through activating PGC-1α and phosphorylating class IIa HDACs to trigger their export from the nucleus, leading to increased mitochondrial biogenesis and fatty acid oxidation^3,12^. However, AMPK activation also limits muscle hypertrophy, growth, and protein synthesis by suppressing mTORC1 and promoting catabolism^13^. Moreover, the enzymatic activities of class IIa HDACs depend on the associated class I enzyme HDAC3^14^, while we found that skeletal muscle-specific deletion of HDAC3 increases susceptibility to diabetes^15^. The balance between the beneficial and detrimental effects of exercise-induced activation of mitogen-activated protein kinases (MAPKs) and nuclear factor-kappa B (NF-kB) remains controversial^16–18^. Similarly, the roles of exercise-induced secreted factors, such as irisin and IL-6, are also debated^19–22^. In summary, the mechanisms underlying exercise-mediated benefits remain incompletely understood.

Exercise-induced production of reactive oxygen species (ROS) is implicated in the benefits of exercise^23^. ROS activates AMPK and calcium/calmodulin-dependent protein kinase (CaMK), which activates PGC-1α, leading to increased mitochondrial gene expression, enhanced mitochondrial biogenesis, and oxidative metabolism^3,4^. ROS can activate or inhibit NF-κB signaling through upstream kinases (IKK and AKT), affecting I-κB degradation, or modifying NF-κB heterodimers to influence nuclear translocation and DNA binding, regulating genes related to inflammation and stress response^24^. Supplementation with antioxidant vitamins C or E during exercise training inhibits improvements in insulin sensitivity as well as the induction of genes involved in mitochondrial biogenesis and antioxidant defense^25,26^. ROS is known to have a biphasic role in muscle force generation and fatigue resistance^27^. Although high-level ROS is generally detrimental, quenching mitochondrial ROS in myofibers during eccentric exercise exacerbated myofiber damage and increased force loss^28^. Likewise, a deficiency in NADPH oxidases, a source of ROS, impaired exercise-induced beneficial adaptations in skeletal muscles^29–31^. The biphasic effects of ROS during exercise exemplify the concept of hormesis, in which low-dose stressors exert beneficial effects. Most prior research has focused on the ’dosage’ of ROS. Beyond dosage, the temporal dynamics of ROS-induced stress responses represent a relatively underexplored dimension.

Exercise-induced beneficial effects on systemic metabolism can be independent of its impact on energy expenditure or body weight loss^32^, and may require transcriptional regulation^33^. Although exercise-induced epigenomic changes have been profiled^34–38^, identifying upstream signaling pathways and their specific transcription factors remains a major challenge. Gene transcription is initiated by enhancers looping with transcription start sites (TSS) or immediate promoters, followed by recruitment of chromatin remodelers and transcription initiation complexes containing RNA polymerase II (Pol2)^39^. Activated Pol2 transcribes DNA in the vicinity of looping complexes in a divergent manner, generating RNAs from both gene bodies and enhancers. While transcripts from gene bodies undergo productive elongation, bidirectional transcripts from the enhancers (eRNA) are short and transient, but are detectable by global nuclear run-on assay (GRO-seq)^40^. The eRNA is a superior marker for active enhancers compared to other enhancer markers: (1) Binding-based profiling, such as chromatin immunoprecipitation (ChIP), does not distinguish functional binding vs. non-functional binding because of the frequent ’on and off’ sampling events during protein-DNA binding^41–43^. (2) Chromatin accessibility assays, such as assay for transposase-accessible chromatin using sequencing (ATAC-seq), DNase I hypersensitivity (DNase-seq), or micrococcal nuclease assays (MNase-seq), have limited ability to distinguish cell lineage–associated hotspots from signaling-specific enhancers. (3) Self-transcribing active regulatory region sequencing (STARR-seq) can generate false positive results because it is independent of endogenous chromatin architecture and genomic context. (4) Compared to Pol2 or p300 ChIP-seq, eRNA has higher dynamic ranges and can mark enhancers in introns due to their bidirectional nature. (5) Compared to histone markers and DNA methylation, eRNAs are small and can locate enhancers more accurately. Thus, GRO-seq provides a powerful non-biased approach to identify exercise-responsive enhancers. No comprehensive profiling of eRNAs in response to exercise has yet been performed.

Here, we utilized GRO-seq to map enhancer activity and RNA-seq to profile transcriptomic changes in adult male C57BL/6 mice immediately following a bout of treadmill exercise. Our data revealed that exercise primarily activates gene transcription at this early phase. Motif analysis of exercise-activated enhancers identified AP-1 as a critical initiator of transcriptomic changes in skeletal muscles. CRISPR/Cas9-mediated screening further suggests that cFos/JunD contributes to contraction-induced transcriptional changes in myotubes.

AP-1 is a well-known stress response signaling, with cFos being the best-characterized immediate early gene (IEG) activated by various stress stimuli, promoting cell proliferation and growth^44^. Intriguingly, AP-1 is also a pioneer factor for senescence^45^ and aging-related chromatin opening^46^. We showed that intermittent but not continuous activation of AP-1 recapitulates exercise benefits, including enhanced contractile performance, glucose tolerance, and mitochondrial volume density. Conversely, disrupting AP-1 dynamics blunts training adaptations. These findings provide a mechanistic explanation for the necessity of recovery cycles in exercise training and highlight the importance of stress-response pulsatility for exercise-mediated benefits. The paradoxical pro- and anti-aging roles of AP-1 further underscore the kinetics-dependent impact of stress and stress responses on senescence and healthspan regulation.

## RESULTS

### Genome-wide analysis of exercise-induced enhancers pinpoint AP-1

To identify exercise-induced enhancers in skeletal muscle, we performed GRO-seq and RNA-seq analyses of the quadriceps muscles of adult male C57BL/6 mice following three consecutive treadmill exercise sessions at 10 m/min. Each exercise session lasted one hour and was followed by a 30-minute rest interval. Tissues were harvested immediately after the last exercise session. RNA-seq identified that most differentially expressed genes (DEGs, exercise vs. rest) were upregulated upon exercise (**Fig. 1A** and **Supplementary Table S1**). These DEGs were highly enriched in glucose metabolism, lipid metabolism, ROS metabolism, nutrient-sensing signaling pathways, unfolded protein stress response, and autophagy (**Fig. 1B-D**). The direction of gene expression changes closely correlated with nascent RNA transcription in gene bodies, suggesting that transcriptional activation underlies these changes (**Fig. 1E**). Motif analysis of the exercise-induced eRNAs identified ’TGA(C/G)TCA’ as the top motif (**Fig. 1F**). This motif is known as the 12-O-Tetradecanoylphorbol-13-acetate (TPA) response element and is recognized by the AP-1 family of transcription factors (TF)^47^. Many exercise-activated genes are accompanied by exercise-activated eRNAs that contain an AP-1 motif (**Fig. 1G**). Although we pooled muscle samples from multiple mice for a single GRO-seq library in this analysis, we also performed RNA-seq and GRO-seq analyses in a mouse line with a skeletal muscle-specific knockout of histone deacetylase 3 (HDAC3) as a biological replicate. HDAC3-depleted muscles were very similar to WT mice in their responses to acute treadmill exercise in transcriptomic changes, correlation between GRO-seq and RNA-seq, and the top-enriched motif in exercise-induced enhancers (**Supplementary Fig. S1** and **Table S2**). These results demonstrate the robustness and reproducibility of the analysis and suggest that exercise-induced AP-1 activation is independent of HDAC3. In summary, unbiased GRO-seq and RNA-seq analyses suggest that AP-1 may be a critical initiator of exercise-induced transcriptomic changes in skeletal muscle.

**Figure 1.**
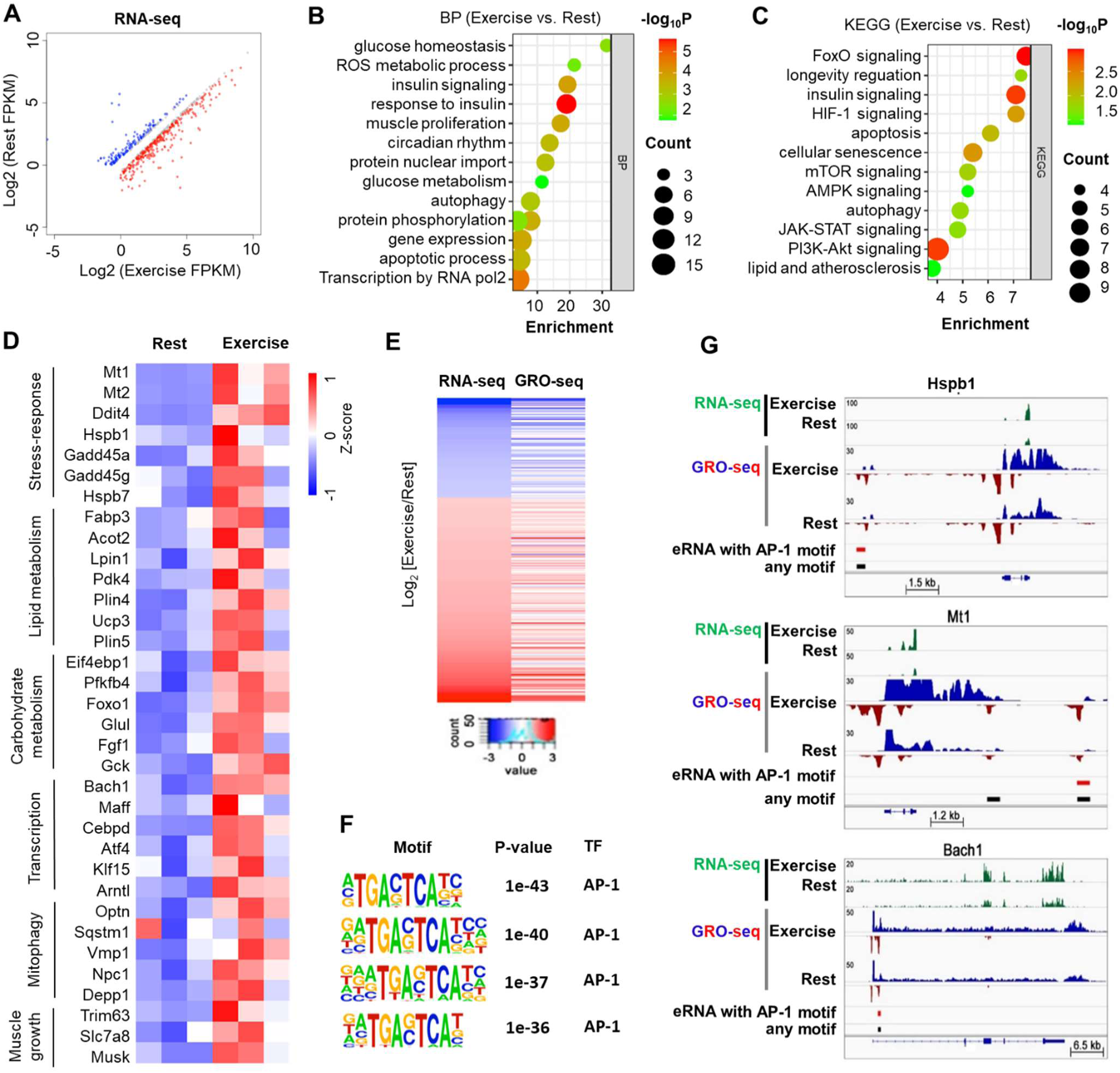
GRO-seq/RNA-seq analysis of exercise-induced enhancers pinpoints AP-1. **(A)** Scatter plot of fragments per kilobase of exon per million fragments (FPKM, log_2_ scale) from RNA-seq of quadriceps muscle in 4-month-old wild-type (WT) male mice comparing exercise vs. rest (n=3, q<0.05, |log_2_(fold-change)| > 0.58). **(B-C)** Gene ontology of biological processes (BP) and Kyoto Encyclopedia of Genes and Genomes (KEGG) pathway enrichment of exercise-induced upregulated DEGs. **(D)** The heat map of the top DEGs shows Z-scores calculated from FPKM. **(E)** Heat maps of RNA-seq and GRO-seq signals at gene bodies of DEGs (reads per million normalized to the bin size). **(F)** Top enriched motifs in exercise-induced eRNAs from WT mice. **(G)** Representative exercise-induced genes with nearby eRNAs containing the AP-1 motif. Y-axis values are reads per million, normalized to the bin size.

### cFos/JunD contributes to contraction-induced transcriptional and metabolic responses

The AP-1 family of basic leucine zipper (bZIP) TFs includes c-Jun, JunB, JunD, cFos, FosB, Jdp2, Fra2, and others, which form heterodimers to bind DNA. To identify which members of the AP-1 family play a predominant role in exercise-induced transcriptional changes, we subjected differentiated C2C12 myotubes to electric pulse stimulation (EPS) as an *in vitro* myofiber contraction model (**Fig. 2A**), which was shown to recapitulate some of the molecular changes in muscles *in vivo* during exercise^48^. EPS at 11 V for 6 hours with a 2-ms pulse duration and 1 Hz frequency led to observable myotube contraction under the microscope and triggered transcriptional activation of genes related to unfolded protein stress and metabolism in differentiated myotubes (**Fig. 2B**) but not in undifferentiated myoblasts (**Supplementary Fig. S2A**). Cell viability assay ruled out the possibility of abnormal myotube death due to EPS (**Supplementary Fig. S2B**). These results validated the EPS-treated myotubes as a useful *in vitro* model to study the transcriptional effects of muscle contraction. We then sought to identify the key AP-1 family members involved in exercise-induced transcriptional regulation. Using adenovirus-mediated CRISPR/Cas9 genome-editing, we individually knocked down each AP-1 family member using two sgRNAs per gene (**Supplementary Table S2**). We used western blot analyses to validate some of the sgRNA-mediated knockdown (**Supplementary Fig. S2C**). Knockdown of cFos or JunD produced the most prominent effects on EPS-induced transcription, suggesting these two AP-1 family members likely play a predominant role in contraction-induced transcriptional activation (**Fig. 2C**).

**Figure 2.**
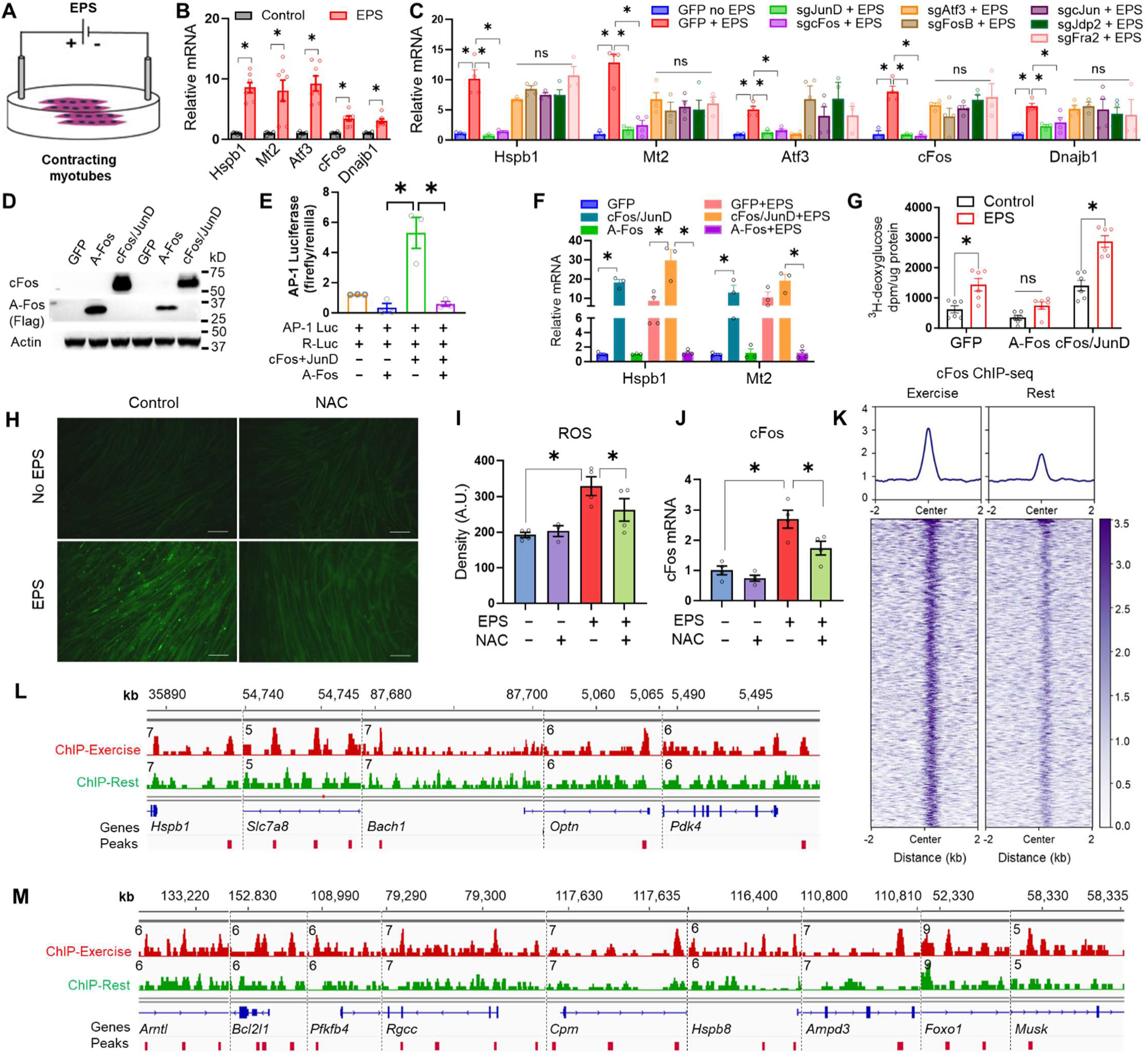
cFos/JunD contributes to contraction-induced transcriptional and metabolic responses *in vitro*. **(A)** Differentiated C2C12 myotubes were subjected to electric pulse stimulation (EPS). **(B)** RT-qPCR analysis of EPS-treated C2C12 myotubes. *p<0.05 by student’s t-test (n = 7–9). **(C)** RT-qPCR of myotubes infected with lentivirus expressing single-guide RNAs (sgRNAs) targeting AP-1 family members, *p<0.05 by 2-way ANOVA and Bonferroni test (n = 4). (**D**) Immunoblot showing tetracycline-inducible cFos/JunD expression and dominant-negative Flag-tagged A-Fos expression in myotubes. (**E**) Luciferase activity from an AP-1-controlled luciferase reporter, normalized to Renilla luciferase in 293T cells transfected as indicated. *p<0.05 by 2-way ANOVA and Bonferroni test, n = 3. (**F**) RT-qPCR of myotubes after adenoviral infection as indicated. * p<0.05 by 2-way ANOVA and Bonferroni test, n = 4. **(G)** Glucose uptake assay with ^3^H-deoxyglucose in myotubes after the indicated adenoviral infection. *p<0.05 by t-test. n = 6. (**H-I**) Fluorescence microscopy of intracellular ROS levels in myotubes with or without the ROS-scavenging agent NAC. Scale bar, 10 μm. A.U., arbitrary unit. **(J)** RT-qPCR of cFos expression. *p<0.05 by t-test. n = 4. (**K**) Heatmap of upregulated 963 cFos ChIP-seq peaks (Exercise vs. Rest), with average read density, from quadriceps muscles of 5-month-old male mice immediately after two bouts of treadmill exercise (10m/min, 1h each) separated by a 30-minute rest interval. **(L-M)** Genome browser tracks of cFos ChIP-seq signals in mouse quadriceps. Data are mean ± SEM.

To explore the functional consequences of AP-1 activation, we engineered adenovirus vectors to manipulate AP-1 activity in differentiated C2C12 myotubes. For gain-of-function (GOF) experiments, cFos and JunD were co-overexpressed under the promoter containing the tetracycline response element (TRE) for inducible expression. For loss of function (LOF), we overexpressed a dominant-negative truncated form of cFos, termed A-Fos, which is known to inhibit the AP-1 binding^49^. Efficient expression was validated by western blot (**Fig. 2D**). Overexpression of cFos/JunD activated AP-1-dependent transcriptional activity in a luciferase reporter gene assay, which was abolished by A-Fos co-overexpression (**Fig. 2E**), validating the GOF and LOF constructs. Adenovirus-mediated AP-1 GOF in differentiated C2C12 cells in the absence of EPS mimicked EPS-induced transcriptional activation for Hspb1 and Mt2, whereas AP-1 LOF abolished EPS-induced upregulation of these genes (**Fig. 2F**). In addition to gene expression analysis, we also sought to investigate the physiological impact of AP-1 activation on glucose uptake using isotope tracer analysis. We found that EPS-induced glucose uptake was impaired upon AP-1 LOF and was mimicked by AP-1 GOF (**Fig. 2G**).

To investigate the upstream mechanism of exercise-induced AP-1 activation, we examined ROS production, since muscle contractions stimulate ROS generation in myofibers, which triggers redox-dependent adaptations^23^. We measured ROS in EPS-challenged C2C12 cells with a ROS probe H2DCFDA (2’,7’-dichlorofluorescein diacetate)^50^ and found an increased cellular ROS level in contracting myotubes (**Fig. 2H-I**). *N*-acetylcysteine (NAC), a ROS neutralizer^51^, blocked EPS-induced ROS accumulation and transcriptional activation of cFos (**Fig. 2H-J**), suggesting that contraction-induced ROS could be an upstream mediator of AP-1 activation. ChIP-seq with cFos antibodies in muscles further identified upregulated cFos binding on genomic loci after a single bolus exercise in mice (**Fig. 2K** and **Supplementary Fig. S3**). Many of these exercise-induced cFos peaks were located near genes that were upregulated by exercise (**Fig. 2L-M**). In summary, these *in vitro* and *in vivo* findings suggest that AP-1 is activated in skeletal muscle after exercise, likely through a ROS-dependent mechanism, and contributes to the transcriptional and metabolic adaptations to muscle contraction.

### Neonate-onset AP-1 loss-of-function (LOF) impairs muscle performance and glucose tolerance

To determine the role of AP-1 in muscle contractile and metabolic functions, we injected neonatal mice carrying skeletal muscle-specific myosin light chain 1f-Cre (MLC-Cre)^52^ with an adeno-associated virus (AAV) vector AAV9.CAG.FLEX.A-Fos expressing A-Fos in a Cre-dependent manner using the FLEX system^53^. Compared to adult administration, subcutaneous injection of AAV in neonates requires a lower viral dose, likely due to the small body size, incomplete connective tissue deposition, and incomplete development of the liver^54,55^. Consistent with previous reports^54–61^, this AAV-dependent approach leads to transgene expression specifically in the skeletal muscle (**Supplementary Fig. S4**). The expression of A-Fos in the tibialis anterior (TA) muscles was confirmed by immunoblot (**Fig. 3A**). In comparison to the wild-type control (Con) mice injected with an empty AAV vector, A-Fos mice showed normal body weight (**Fig. 3B**), body length (**Fig. 3C**), body composition (**Fig. 3D**), and food intake (**Fig. 3E**). However, stride, stance, and sway were significantly reduced in A-Fos mice compared to the control (**Fig. 3F-I**), indicating a disruption in motor function. Indeed, A-Fos mice demonstrated compromise in the vertical wire hanging test (**Fig. 3J**), the horizontal grip strength test (**Fig. 3K**), and the graded treadmill exercise test (**Fig. 3L-M**). These functional impairments were not associated with changes in muscle mass (**Fig. 3N**), fiber-type composition (**Fig. 3O-P**), or cross-sectional fiber diameter (**Fig. 3Q**), but were accompanied by reduced glucose tolerance (**Fig. 3R-S**). These results suggest that muscle-specific AP-1 LOF from the neonatal stage impairs muscle contractile performance and disrupts glucose homeostasis without affecting muscle mass or fiber-type composition.

**Figure 3.**
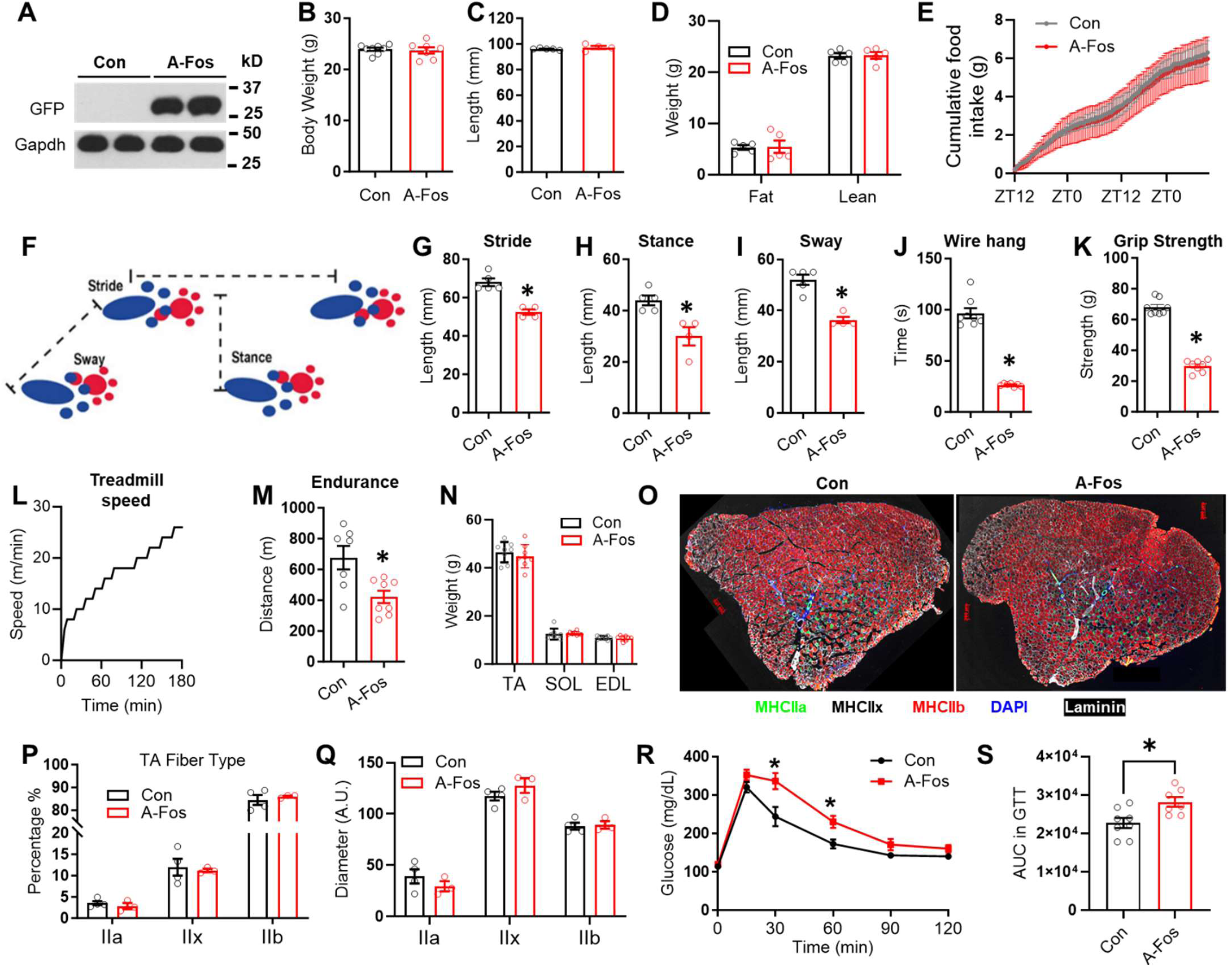
Neonate-onset AP-1 loss-of-function impairs muscle performance and glucose tolerance. **(A)** Immunoblot of tibialis anterior (TA) muscles from 6-month-old adult mice after neonatal subcutaneous injection of AAV9.FLEX.A-Fos, a dominant-negative truncation of cFos (A-Fos group), or AAV9.FLEX.GFP control (Con). **(B-D)** Body weight, body size, and body composition by magnetic resonance imaging (MRI) at 4 months old. **(E)** Food intake. **(F)** Illustration of gait parameters, including stride length, stance width, and sway distance. **(G–I)** Quantification of stride, stance, and sway at 4 months. **(J-K)** Wire hang test and grip strength at 4 months old. **(L-M)** Treadmill speed profile and endurance test at 6 months. *p<0.05 by 2-way ANOVA and Bonferroni’s test, n = 7-8 mice. **(N)** Muscle mass at 6 months old. TA, tibialis anterior; SOL, soleus; EDL, extensor digitorum longus. **(O-Q)** Immunofluorescence staining of TA muscles from control and A-Fos mice with antibodies or reagents against myosin heavy chain (MHC) isoforms: MHCIIa (green), MHCIIx (black), MHCIIb (red), DAPI (blue), and laminin (white). n = 4–5 mice. **(R-S)** Glucose tolerance test and the area under the curve (AUC) of the glucose excursion in 3-month-old adult mice. n = 7-8 mice. *p<0.05 by 2-way ANOVA and Bonferroni’s test. Data are mean ± SEM.

### Adult AP-1 LOF blunts exercise-induced improvement in muscle performance but not glucose tolerance

Since A-Fos overexpression from the neonatal stage altered muscle contractile function at the non-training baseline, we were unable to test the role of AP-1 in exercise training-induced effects. We suspect that neonate-onset A-Fos overexpression might affect postnatal muscle development. To bypass this potential confounding effect, we engineered dual AAV vectors for adult-onset, inducible A-Fos (iA-Fos) using the Tet-On system^62^. A Flag-tagged A-Fos is driven by the TRE-tight promoter that is inducible by doxycycline (DOX) in the presence of rtTAM2. A second AAV vector expresses rtTAM2 in a Cre-specific manner (**Fig. 4A**). Thus, administration of the dual AAV vector into MLC-Cre mice leads to DOX-inducible, muscle-specific A-Fos expression. A-Fos was efficiently induced in the muscles after DOX treatment via drinking water in adult mice **(Fig. 4B)**. Compared to the WT control mice (Con) that received only one AAV vector and were similarly treated with DOX, iA-Fos mice exhibited normal voluntary wheel-running activity (**Supplementary Fig. S5A**), normal body weight **(Fig. 4C**), and normal body length (**Supplementary Fig. S5B**). Importantly, iA-Fos mice showed normal stride, stance, and sway at the baseline **(Fig. 4D-F)**, in contrast to the neonate-onset A-Fos overexpression. *Ex vivo* muscle physiology analysis showed no differences in basal muscle force and fatigue resistance in the iA-Fos mice (**Supplementary Fig. S5C-E**). These findings suggest that iA-Fos did not impact baseline muscle contractile function, allowing us to address the specific role of AP-1 in exercise training adaptations. Toward this goal, we subjected mice to voluntary training on running wheels for 3 months, which led to a moderate improvement in grip strength **(Fig. 4G)** and a robust improvement in the graded treadmill exercise test in the WT control mice (**Fig. 4H-I**). These training-induced improvements were blunted in mice with iA-Fos expression (**Fig. 4H-I**), demonstrating that muscle AP-1 is required for training-induced adaptations in contractile performance and glucose metabolism.

**Figure 4.**
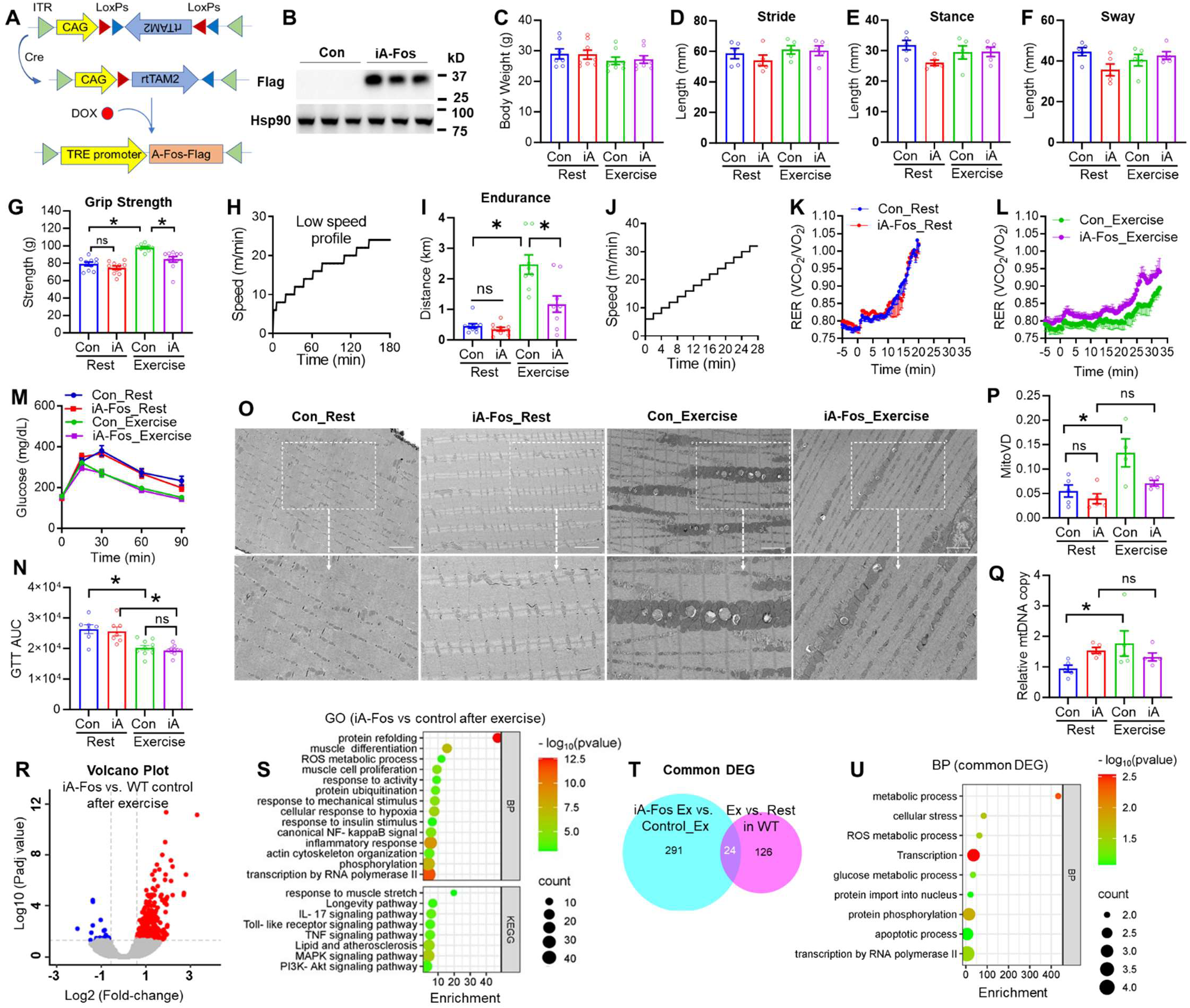
Adult-onset AP-1 loss-of-function blunts exercise-induced benefits in muscle performance, but not glucose tolerance. **(A)** Schematic for the inducible, muscle-specific A-Fos overexpression. **(B)** Immunoblot of TA muscles from mice with inducible A-Fos (iA-Fos) overexpression after doxycycline (DOX) treatment. Mice injected with rtTA/GFP and treated with DOX served as the control (Con). **(C-F)** Body weight and gait analysis of stride, stance sway distance at 3 months of age. **(G)** Grip strength at 7 months old, n = 9-10 mice per group. **(H-I)** Treadmill speed profile and endurance test at 5 months of age. *p<0.05 by 2-way ANOVA and Bonferroni’s test, n = 8-10 mice per group. **(J-M)** Treadmill speed profile and respiratory exchange ratio (RER) at 9 months old, n = 4-7 mice. **(M-N)** Glucose tolerance test and area under the curves (AUC) at 4 months old, n = 7-10 mice. iA, iA-Fos. *p<0.05 by 2-way ANOVA and Bonferroni’s test. **(O)** Electron microscopy of TA muscle. **(P)** Quantification of mitochondrial volume density (MitoVD) from 4-5 randomly captured images. *p<0.05 by 2-way ANOVA and Bonferroni’s test. MitoVD was calculated as the mitochondrial area (MA) divided by the total area (TOA). Scale bar, 2 µm. **(Q)** qPCR of mitochondrial DNA (mtDNA) copy number. *p<0.05 by 2-way ANOVA and Bonferroni’s test. n = 5 mice. **(R)** Volcano plot of differentially expressed genes (DEGs, q<0.05, |log_2_(fold-change)|>0.58) between WT and iA-Fos muscles after a bout of exercise. **(S)** Bubble plot of pathway enrichment of DEGs (iA-Fos vs. WT). **(T)** Venn diagram of iA-Fos-induced DEGs (iA-Fos vs. control) and exercise-induced DEGs (Exercise vs. Rest). **(U)** Bubble plot of pathway enrichment of common DEGs. Data are mean ± SEM.

The voluntary wheel-running exercise training did not change muscle mass in either wild-type control or iA-Fos mice (**Supplementary Fig. S5F**). We used indirect calorimetry to assess fuel metabolism during exercise. Non-trained iA-Fos mice showed a normal intensity-dependent increase in the respiratory exchange ratio (RER) during a treadmill test with increasing speed (**Fig. 4J**). However, after training, iA-Fos mice showed a faster increase in RER than the WT control mice (**Fig. 4K**). Since a higher RER indicates greater reliance on carbohydrates and is a sign of exhaustion, this data is consistent with the compromised endurance improvement (**Fig. 4J-L**). The higher RER in iA-Fos mice could be due to an intrinsic fuel preference toward glucose or against fatty acids. However, iA-Fos mice showed a similar degree of improvement in glucose tolerance as wild-type control mice after chronic exercise training (**Fig. 4M-N**). Isotope-based glucose uptake assay in freshly isolated muscles showed that exercise-induced glucose uptake was similar between iA-Fos mice and WT control (**Supplementary Fig. S5G-H**). Thus, the high RER is likely due to compromised muscle endurance and is not driven by an intrinsic change in glucose metabolism.

Since mitochondria play a critical role in muscle bioenergetics, we sought to characterize mitochondrial adaptations to exercise. Mitochondria undergo dynamic changes in response to exercise, including repair, biogenesis, size modulation, turnover, fusion, and fission, to meet increased energy demands during physical activity. Transmission electron microscopy revealed that exercise increases mitochondrial volume density (mitoVD) in WT control mice, whereas iA-Fos expression blocked such changes (**Fig. 4O-P**). iA-Fos also blocked training-induced changes in mitochondrial DNA (mtDNA) copy number (**Fig. 4Q**), while no significant differences were found in oxidative phosphorylation (OXPHOS) protein levels among groups (**Supplementary Fig. S5I**). The exercise-induced changes in WT mice are consistent with several previous studies reporting an exercise-induced increase in mitoVD in both human and rodent skeletal muscle^63–65^, although other studies reported various effects of exercise on mitochondrial properties^66,67^. These results indicate that mitochondrial adaptations can vary based on exercise intensity, duration, muscle fiber type, and species. Nonetheless, since iA-Fos blocked the exercise training-induced mitochondrial changes in our study, it demonstrates that muscular AP-1 activation is required for training-dependent mitochondrial adaptations.

RNA-seq analysis comparing iA-Fos vs. WT muscles harvested after acute treadmill running exercise identified 315 differentially expressed genes (DEGs) enriched in stress response, protein folding, muscle differentiation, ROS metabolic processes, and insulin signaling (**Fig. 4R-S** and **Supplementary Table S3**). Overlapping these DEGs with previously identified transcriptomic effects of exercise (acute exercise vs. rest in wild-type mice) identified 24 overlapping DEGs (**Fig. 4T**) that are enriched in cellular stress response and metabolic processes (**Fig. 4U**). Contrary to the expectation that AP-1 LOF would compromise exercise-induced transcriptomic changes, most of these DEGs were upregulated, instead of downregulated, in iA-Fos vs. WT control. We speculate that AP-1 LOF could delay the resolution of exercise-induced stress, leading to the buildup of stress and activation of other stress-responsive signaling pathways. Thus, timely activation of AP-1 to quickly restore cellular homeostasis could be critical for exercise-induced hormesis. These results demonstrate that AP-1 activation is required for normal transcriptomic responses to exercise and exercise-mediated improvements in muscle contractile performance and mitochondrial dynamics.

### AAV-mediated intermittent, but not continuous, AP-1 activation recapitulates exercise benefits

To explore whether AP-1 activation can mimic exercise benefits, we engineered AAV vectors with the Tet-ON system for inducible AP-1 overexpression. We used a Cre-dependent FLEX system to restrict rtTA to skeletal muscle in one AAV vector (AAV9.CAG.FLEX.rtTAM2.T2A.GFP), and the TRE promoter to drive the DOX-inducible expression of cFos and JunD in another AAV vector (AAV9.TRE9.JunD.T2A.cFos) (**Fig. 5A**). We first validated the vectors in cultured HEK293 cells after transfection (**Supplementary Fig. S6A-B**). After subcutaneous neonatal injection of the dual AAV vectors into the muscle-specific MLC-Cre mice, we introduced doxycycline (DOX) to adult mice in drinking water for 3 days, followed by a washout for 4 days (3d-ON, 4d-OFF) to establish intermittent AP-1 overexpression (iOE). We reason that such intermittent overexpression would mimic, to a certain degree, exercise-induced temporary AP-1 activation followed by stress resolution. We also included a group with continuous AP-1 overexpression (cOE) by maintaining DOX in the drinking water throughout. Mice receiving empty AAV vectors and similar DOX administration served as respective wild-type (WT) control groups (iCon or cCon). The expression of AP-1 in the muscles of iOE mice was confirmed through immunoblot (**Fig. 5B**). Compared to its control, iOE mice showed normal body composition (**Fig. 5C**), oxygen consumption (**Fig. 5D**), and food intake (**Supplementary Fig. S6C**). However, iOE mice exhibited a longer duration in the wire-hang test compared to their control (**Fig. 5E**), while cOE mice did not show such improvement (**Fig. 5E**). The iOE mice, but not cOE mice, showed a modest improvement in grip strength (**Fig. 5F**) and treadmill test (**Fig. 5G-H**). No significant change in muscle mass was observed in iOE mice or cOE mice compared to their controls (**Fig. 5I**). These results suggest that intermittent, but not continuous, AP-1 activation improves muscle contractile performance *in vivo*.

**Figure 5.**
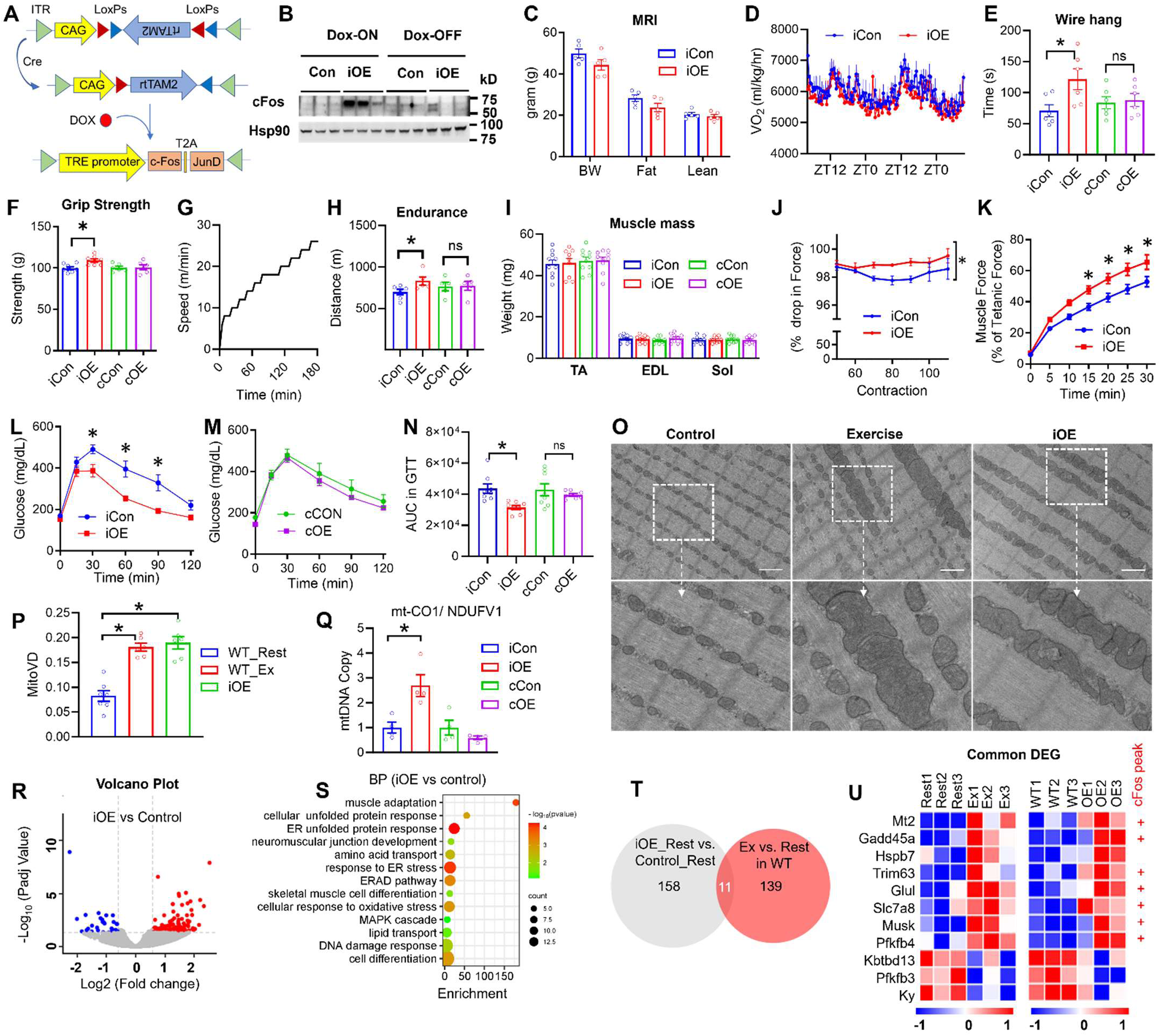
Intermittent, but not continuous, overexpression of AP-1 in muscles improves contractile performance and glucose tolerance. **(A)** Schematic of the Tet-ON and FLEX systems for muscle-specific, inducible AP-1 overexpression. **(B)** Immunoblot of EDL muscle from 11-month-old mice with inducible overexpression (iOE) of cFos/JunD after doxycycline (DOX) treatment. **(C)** Body weight (BW), lean, and fat mass by magnetic resonance imaging (MRI) at 11 months of age. n = 4 mice. **(D)** Oxygen consumption (VO_2_) at 11 months old, n = 3-4 mice. **(E-F)** Wire hang test and grip strength test at 4 months old, n = 5-8 mice. **(G-H)** Treadmill speed profile and endurance test at 4 months old, n = 5-6 mice. **(I)** Muscle weight. (J-K) *Ex vivo* muscle contraction physiology analysis of EDL muscles at 11 months old, n = 4 mice. **(L-N)** Glucose tolerance test and the area under the curves (AUC) at 7 months old, n = 8 mice. **(O-P)** Electron microscopy images and quantification of mitochondrial volume density (MitoVD) of TA muscle at 5 months old. Scale bar, 1 μm. Image quantification was performed using Qpath with blinded acquisition from 4-5 randomly captured images. **(Q)** qPCR analysis of mitochondrial DNA copy number from EDL muscles at 9 months old, n = 4 mice. **(R)** Volcano plot of RNA-seq analysis of EDL muscles comparing iOE with WT control. **(S)** Bubble plot of GO and KEGG pathway enrichment analysis of DEGs (iOE vs. WT, q<0.05, |log_2_(fold-change)|>0.58). **(T)** Venn diagram comparing iOE-activated DEGs (iOE vs WT) and exercise-activated DEGs (exercised vs. rest in WT). **(U)** Heatmap of the genes activated by both iOE and exercise, with cFos binding sites. Data are mean ± SEM.

Consistent with the *in vivo* results, an *ex vivo* physiology test using isolated EDL muscles of iOE mice displayed a slower rate of force dropping (**Fig. 5J**) and faster recovery from fatigue (**Fig. 5K**) compared to the WT control during repeated exhaustive contractions, demonstrating muscle-autonomous improvement in contractile performance. iOE mice also showed improved glucose tolerance compared to the WT control, whereas cOE mice did not show this improvement (**Fig. 5L-N**). These data suggest that intermittent AP-1 activation positively affects metabolic health, mimicking the benefits of regular exercise. Electron microscopy analysis showed that intermittent AP-1 overexpression increased mitochondrial volume density (**Fig. 5O-P**) and mitochondrial DNA copy number (**Fig. 5Q**). Continuous overexpression of AP-1 did not alter any of these outcomes. Thus, intermittent AP-1 activation can mimic some exercise benefits, including enhanced muscle performance and metabolic improvements, whereas continuous AP-1 activation fails to achieve these outcomes.

RNA-seq of the TA muscles identified 169 DEGs in iOE mice compared to WT control, which are enriched in cellular stress response, muscle differentiation, and metabolism (**Fig. 5R-S** and **Supplementary Table S4**). After overlapping these iOE-induced DEGs with the exercise-induced DEGs (acute exercise vs. rest in wild-type mice), we found 11 overlapping DEGs. Eight of these common genes are upregulated by both exercise and iOE compared to their respective controls and have cFos binding peak nearby (**Fig. 5T-U**). These common DEGs are involved in stress response and glucose metabolism. These results indicate intermittent AP-1 overexpression partially recapitulates exercise-induced transcriptomic changes and improves contractile performance, glucose metabolism, and mitochondrial volume density.

### Intermittent, but not continuous, transgenic AP-1 activation recapitulates exercise benefits

AAV-mediated AP-1 overexpression could be confounded by virus-induced toxicity. Therefore, we constructed an AP-1 transgenic mouse line expressing TRE.JunD.2A.cFos and crossbred it with the Rosa26-LSL-rtTA3 (JAX #029617) and MLC-Cre mice (**Supplementary Fig. S7**). The resultant TRE-JunD.2A.cFos/rtTA3/MLC-Cre mouse line allows for DOX-inducible expression of AP-1 (cFos/JunD) specifically in skeletal muscles. We used the 3d-ON/4d-OFF regimen of DOX treatment in the drinking water to induce intermittent AP-1 overexpression (iAP-1) and validated it by immunoblot (**Fig. 6A**). We set up another group by continuously maintaining DOX in the drinking water and named it continuous AP-1 overexpression (cAP-1). The WT control mice carrying rtTA3/MLC-Cre or TRE-JunD.2A.cFos/MLC-Cre were treated with DOX intermittently or continuously, and were referred to as iCon or cCon, respectively. Both iAP-1 and cAP-1 showed normal body weight compared to their respective controls (**Fig. 6B**). Compared to iCon, iAP-1 mice showed improved wire-hang performance (**Fig. 6C**), a modest increase in grip strength (**Fig. 6D**), and a robust improvement in the graded treadmill exercise test (**Fig. 6E-F**). In contrast, cAP-1 mice did not exhibit such improvements in any of the tests (**Fig. 6C-F**). Neither iAP-1 nor cAP-1 mice exhibited changes in muscle mass compared to their respective controls (**Fig. 6G**), suggesting that the observed functional changes were not due to muscle hypertrophy. *In vivo* analysis of dorsiflexor muscle contractile physiology revealed that iAP-1 mice, but not cAP-1 mice, displayed enhanced muscle strength (Fig. 6H) and fatigue resistance (Fig. 6I-L) compared with their respective controls. Consistently, glucose tolerance was improved in iAP-1 mice but not cAP-1 mice (**Fig. 6M-N**). In line with the glucose tolerance test, isotope tracing studies showed that glucose uptake is upregulated in iAP-1 mice but not cAP-1 mice, compared to their respective controls (**Fig. 6O**). Consistently, iAP-1 muscles showed increased mitochondrial DNA copy number compared to their WT control, while cAP-1 did not (**Fig. 6P**), without altering the OXPHOS protein levels (**Supplementary Fig. S7**).

**Figure 6.**
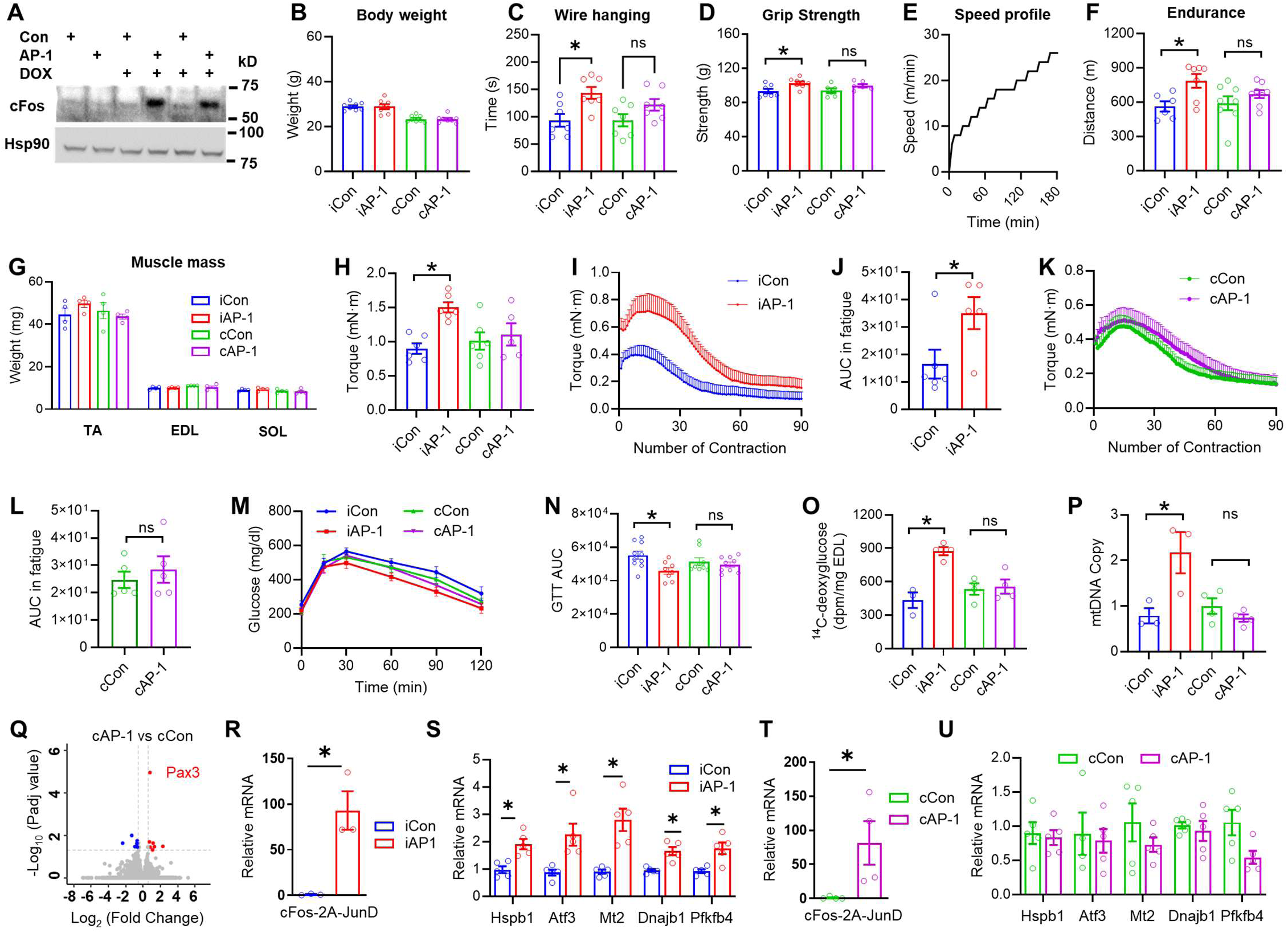
Intermittent, but not continuous, transgenic AP-1 activation recapitulates exercise benefits in contractile performance and glucose tolerance. **(A)** Immunoblot of TA muscles from mice carrying all three alleles (MLC-Cre/rtTA3/AP-1) treated with doxycycline (DOX) for 4 months in an intermittent pattern (iAP-1) or continuous pattern (cAP-1). The control mice carried 2 of the 3 alleles and were treated with DOX similarly. **(B-D)** Body weight, wire hang test, and grip strength. **(E-F)** Treadmill speed profile and endurance test measurement in Con_ iAP-1 (control for iAP-1), iAP-1, Con_cAP-1 (control for cAP-1), and cAP-1 (continuous AP-1) mice at 7 months old, n = 6-8. **(G)** Relative muscle mass (g) of TA, EDL, and soleus muscles from intermittent or continuous AP-1 transgenic and control mice. **(H)** In-vivo dorsiflexion tetanic force measured in TA muscles using the Aurora system, at 6 months old, n = 5-6. **(I-L)** In-vivo muscle fatigue and area under the curve (AUC) calculation in TA muscles using the Aurora system, at 6 months old, n = 5-6. **(M-N)** Glucose tolerance and AUC at 7 months age, n = 8-10. **(O)** Glucose uptake in EDL muscle from iAP-1, cAP-1, and control mice at 8 months age, n=3-4. (**P)** RT-qPCR analysis of mtDNA content in EDL muscle, n = 3-4. **(Q)** Volcano plot of RNA-seq from TA muscle of cAP-1 mice, with only 18 DEGs (q < 0.05, |log_2_(fold-change)|>0.58). **(R-U)** RT-qPCR analysis of TA muscle showing the cFos-2A-JunD transgene and downstream genes, n = 4-5, *p<0.05 by 2-way ANOVA and Bonferroni’s test.

In contrast to the 158 DEGs from iOE vs. WT control muscles, RNA-seq analysis of muscle tissue from cAP-1 mice found only 18 DEGs compared to their WT controls. None of the DEGs from cAP-1 vs WT overlapped with the DEGs in WT exercise vs. WT rest (**Fig. 6Q** and **Supplementary Table S5**). RT-qPCR validation in TA muscle confirmed that iAP-1, but not cAP-1, upregulated classical AP-1 target genes associated with the cellular stress response **(Fig. 6R-U)**. Despite the robust expression of the cFos and JunD transgenes in both iAP-1 and cAP-1 conditions **(Fig. 6R** and **6T)**, only iAP-1 muscle exhibited downstream gene activation, suggesting that forced continuous AP-1 overexpression in the cAP-1 group is transcriptionally ineffective. Prolonged AP-1 activation may lead to its own desensitization through negative feedback mechanisms. For example, cFos expression is regulated by rapid protein degradation via the ubiquitin-proteasome pathway^68^, self-repressive feedback on its transcription^69^, and phosphorylation-dependent protein degradation^70^. These mechanisms may underlie the transcriptional inefficiency of the continuous AP-1 overexpression. In summary, the transgenic mouse models validated the AAV-based results and demonstrated that intermittent, but not continuous, AP-1 activation in skeletal muscle mimics exercise training in improving muscle contractile performance and systemic glucose homeostasis.

## DISCUSSION

Our results revealed that AP-1, a pioneer factor for senescence^45^ and aging-related chromatin opening^46^, plays a positive role in adaptive responses to exercise, a lifestyle intervention that counteracts aging and extends lifespan. The role of AP-1 in promoting senescence is supported by the findings that silencing AP-1 suppresses senescence and activates pro-proliferative genes^45^, while overexpressing AP-1 can mimic stress-induced chromatin opening^46^. Blocking AP1 activity extends the lifespan of Drosophila, albeit with increased oxidative damage^71^. Increased AP-1 (cFos/Jun) gene expression is a conserved feature of immune aging and contributes to chronic inflammation^72,73^. The AP-1 member ATF3 is required for the expression of senescence-associated signature genes in fibroblasts^74^. Consistently, ATF3-binding motifs show increased chromatin accessibility in senescent vs. non-senescent cells, and another AP-1 member, JunB, activates senescence-related gene transcription in muscle satellite cells^75^. However, aging-related low telomerase reverse transcriptase expression can be reversed by activating AP-1^76^.

In contrast to its role in promoting senescence and inflammaging, AP-1 is also activated by exercise, an intervention known to counteract aging and extend healthspan. An acute running session caused a significant transient upregulation of cFos and Jun mRNA, protein, or activity^77–80^. TF binding site enrichment analysis of 27 human muscle transcriptomic datasets suggested that acute exercise activates AP-1^81^. However, long-term aerobic exercise downregulates cFos in muscles in an integrated analysis of several datasets^82^. Chronic exercise training attenuates the acute induction of cFos mRNA and AP-1 motif activity following an exercise bout in mouse muscles compared to the untrained controls^83,84^.

Our findings shed light on the paradoxical dynamics of AP-1 in trained muscle, offering a temporal perspective on its divergent roles in promoting senescence versus mediating exercise-induced benefits. Our parallel *in vivo* comparison demonstrated that intermittent, rather than continuous, overexpression of cFos/JunD in skeletal muscle enhances muscle performance, increases mitochondrial volume density, improves glucose tolerance, and alters the expression of genes involved in stress response and metabolism. These findings indicate that the temporal dynamics of AP-1 activity in muscle are critical for exercise-induced adaptations. Our study aligns with the role of AP-1 in muscle regeneration. cFos has been shown to facilitate early activation of muscle satellite cells (MuSCs) following muscle injury and to promote muscle repair^85^. However, persistent cFos activation *in vitro* in primary muscle progenitor cells impairs their differentiation and myotube formation^86^.

We postulate that the seemingly contradictory pro-aging and anti-aging roles of AP-1 reflect the complex interplay between stress and stress responses. The stress response resolves stress, returning the system to baseline homeostasis. Compared to the low-stress baseline associated with prolonged stress responses (**Supplementary Fig. S8A**), intermittent moderate stressors enhance stress response sensitivity and capacity, enabling faster stress resolution when facing a severe stressor, promoting resilience (**Supplementary Fig. S8B**). This adaptation leads to an attenuated stress response to regular stressors, as rapid resolution removes the need for sustained response, consistent with dampened cFos in trained muscles. However, blocking the stress response blunts the beneficial hormetic effects of intermittent moderate stressors (**Supplementary Fig. S8C**), as indicated by our iA-Fos results. Conversely, intermittent activation of the stress response without stressors can mimic the hormetic benefits (**Supplementary Fig. S8D**), as supported by our iOE and iAP-1 studies. The stress response is usually self-limiting in the absence of stressors, as shown in our cAP-1 studies. However, when stress is unresolved, excessive stress can prolong activation of the stress response (**Supplementary Fig. S8E**), leading to chronic inflammation, tissue damage, and aging, consistent with AP-1’s senescence-promoting role. This framework highlights the importance of the timing and dynamics of stressors and stress responses in shaping senescence and healthspan.

Our observation that continuous AP-1 activation *in vivo* failed to generate benefits aligns with the finding that cFos activity is regulated by self-repressive feedback on its transcription^69^, rapid protein degradation via the ubiquitin-proteasome pathway^68^, and phosphorylation-dependent degradation^70^. The refractory period following cFos activation may contribute to the lack of sustained cFos activity under the continuous overexpression condition. By comparison, intermittent AP-1 overexpression could overcome such desensitization and enable more efficient downstream gene activation, mimicking the transient, physiological AP-1 activation during exercise. Downstream target genes of AP-1 may also contribute to the lack of benefit from continuous AP-1 overexpression. Pax3 emerged as a transcription factor strongly upregulated in cAP-1 compared to WT muscle, but not in iAP1 compared to WT muscle. Pax3 gene expression is typically restricted to muscle progenitors and a subset of quiescent satellite cells in adult skeletal muscle. Given that AP-1 transgene expression is confined to mature myofibers via MLC-Cre in our study, the observed Pax3 upregulation suggests either a non-cell-autonomous effect in stem cells or an induction of a progenitor-like status in myonuclei. Although the precise cellular source of Pax3 in this context remains unclear, its elevated expression may indicate a regenerative block or altered progenitor dynamics in cAP1 muscle. For example, Pax3 can recruit coactivators CBP/p300, which are also essential for AP-1-dependent transcription^87,88^, and may antagonize AP-1 activity via coactivator competition^89^. cAP1 downregulates genes associated with cytoskeletal, mitochondrial, and contractile function. Among them is Klf3, a transcriptional repressor that promotes differentiation by antagonizing progenitor genes such as Pax3, which may contribute to the persistent expression of Pax3 in cAP-1 muscles. Given that Klf3 is predominantly expressed in differentiating myoblasts and mature muscle fibers, its downregulation by cAP1 could reflect a failure to exit the progenitor state, reinforcing a regenerative block or stalled differentiation. In summary, we suspect that Pax3 may act as a molecular brake, indirectly limiting AP-1-driven adaptations under continuous AP-1 activation, shifting the balance from productive remodeling in the presence of intermittent AP-1 activation towards a dysfunctional, non-adaptive state. Future studies employing cell-type-resolved transcriptomics will be necessary to determine the cellular origin and mechanistic contributions of Pax3 in this setting.

Several other target genes downstream of AP-1 could regulate adaptations to exercise. For example, iOE upregulates PFKFB4 expression while downregulating PFKFB3. PFKFB3 shunts glucose into glycolysis, facilitating rapid ATP production for immediate energy demands. In contrast, PFKFB4 enhances flux through the pentose phosphate pathway, supporting redox homeostasis and is essential for lipid biosynthesis and detoxification of ROS^90^. Thus, AP-1-mediated modulation of PFKFB3/PFKFB4 could contribute to metabolic reprogramming that facilitates muscle adaptation to exercise by protecting against oxidative damage and optimizing energy utilization for muscle function.

The current study has several limitations. (1) Perhaps the most intriguing finding is that the benefits of AP-1 activation in muscles depend on its intermittent, but not continuous, expression. While we provide evidence that continuous activation leads to transcriptional incompetence, likely through well-established negative feedback mechanisms, the precise chromatin and co-regulator dynamics remain to be fully elucidated. Future work with higher temporal resolution, examining epigenomic changes and protein-complex formation after each pulse of activation, will be needed to decode the exact mechanisms enforcing this requirement for pulsatility. (2) The current study is based on mouse models. Given the differences between mice and humans in their muscle physiology, it is unclear whether pharmacologically tuning AP-1pulsatility represents a novel therapeutic strategy for human subjects unable to exercise. Future studies correlating AP-1 activity in human muscle biopsies with different training modalities will be crucial to bridge this translational gap. (3) We used bulk RNA-seq and GRO-seq. Therefore, the intercellular communication among various cell types is unclear. (4) Our data suggest ROS as an upstream activator of AP-1, primarily based on the inhibitory effect of the broad-acting antioxidant NAC. The use of NAC and a general ROS probe does not identify the specific redox species or its subcellular origin.

Future work employing more targeted tools, such as genetically encoded redox sensors, mitochondria-specific antioxidants, or inhibitors of particular NADPH oxidase isoforms, will be required to pinpoint the precise redox mechanism linking muscle contraction to AP-1 activation. (5) It is unclear how AP-1 interplays with other exercise-responsive signaling pathways within myocytes. Our work positions intermittent AP-1 activation as a critical, parallel pathway co-opted by exercise, without diminishing the established roles of PGC-1α or AMPK. Indeed, it is likely these pathways operate in a coordinated network, as suggested by a previous study^91^. A key open question is how the rapid, transient AP-1 pulse initiates a transcriptional program that may subsequently engage and synergize with regulators like PGC-1α to orchestrate full metabolic and morphological adaptation. Future studies exploring the interactions between AP-1 and these canonical pathways will be essential for building a complete hierarchical model of exercise-induced transcription.

In summary, our findings underscore the importance of intermittent AP-1 activation in promoting adaptive muscle responses to exercise. The ability to manipulate AP-1 activity through genetic and pharmacological means could open new avenues for combating age-related muscle decline. Future interventions could incorporate training modalities that optimize AP-1 signaling, potentially improving outcomes in aging populations and those with metabolic disorders.

## METHODS

### Cell culture, MTT, luciferase

Mouse myoblast C2C12 (ATCC, CRL-1772) and HEK293T cell lines were maintained in Dulbecco’s Modified Eagle’s Medium (DMEM; Gibco 45000-304, 4.5 g/L glucose) supplemented with 1% streptomycin-penicillin (VWR 12001-692), and 10% fetal bovine serum (FBS) (Fisher SH3007903) in a humidified incubator at 37^◦^C with 5% CO2. Induction of C2C12 differentiation was done by switching to supplementation DMEM with 2% horse serum (Fisher 12001-692) for 5-7 days until mature, multinucleated myotubes were formed. To model exercise-induced contraction *in vitro*, differentiated myotubes were subjected to electric pulse stimulation (EPS) with the C-Pace EP Cell Culture Stimulator (ION Optix, CLD12WA) at 11 V for 6 hours in 12-well plates, with 2 ms pulse duration and 1 Hz frequency^48^. To determine the stress caused by EPS, cell viability was assessed using the MTT (3-(4,5-dimethylthiazol-2-yl)-2,5-diphenyltetrazolium bromide) assay. Intracellular reactive oxygen species (ROS) were measured using 2′,7′-dichlorofluorescein diacetate (DCFH-DA), as described previously^92^. Cells were pretreated with 5 uM N-acetylcysteine (NAC; Sigma A9165) for 1 hour to neutralize ROS, followed by treatment with 2 μM DCFH-DA. After incubation, cells were washed three times with culture medium to remove excess DCFH-DA. Fluorescence was evaluated using a Leica microscope (Dmi8 automated), and captured images were processed and analyzed using ImageJ (NIH). To measure AP-1 transcriptional activity, a dual luciferase reporter assay was performed in 24-well plates. Cells were transfected with a firefly luciferase reporter construct containing AP-1 response elements (AP-1-Luc) and a Renilla luciferase (R-Luc) plasmid as an internal control. Cells were co-transfected with expression plasmids for cFos, JunD, or A-Fos. After 24 hours, tetracycline was added to a final concentration of 1 µg/mL (from a 1 mg/mL stock in 70% ethanol), followed by another 24 hours of incubation. Firefly luciferase signal was normalized to renilla luciferase and reported as the ratio of firefly to renilla luminescence (relative luciferase units, RLU).

### Animals

Animal studies were approved by the Institutional Animal Care and Use Committee (IACUC) at Baylor College of Medicine. C57BL/6 mice were housed at 22 ± 2°C, with 60% ± 5% humidity under standard 12-hour light/dark cycles. Mice were exercised on the Columbus Exer 3/6 treadmill with Detection Unit (1055-SDRM-D64) following the protocol^15^ with certain modifications. During the 3-4 days prior to the experiment or harvest, the mice were pre-trained for 10 min every day to adapt to treadmill running and reduce stress. For transcriptomic analysis following an acute bout of exercise, mice underwent three 1-hour treadmill running sessions at 10m/min, separated by 30 minutes of rest. For ChIP-seq analysis, a separate cohort performed two sessions under the same conditions. Muscle tissue was harvested immediately after exercise. The wheel-running experiment was performed by subjecting mice to voluntary wheel running (PT2-MCR1). For doxycycline treatment, doxycycline (Sigma D9891) was dissolved at 0.4-1 mg/ml in drinking water along with 1-2% sucrose and provided ad libitum to adult mice in either an intermittent (3d ON, 4d OFF) or continuous pattern. Dissected tissues were snap-frozen in liquid nitrogen before extraction for RNA, protein, or nuclear isolation.

### Recombinant DNA and virus for in vitro studies

Adenovirus was used to deliver Cas9 and sgRNAs targeting AP-1 members (Atf3, Fos, Jun, JunD, Jdp2, Fra2/Fos2). We employed the pAd/BLOCK-iT™ system (Invitrogen). sgRNAs were driven by the U6 promoter. Target sequences for sgRNAs were designed specifically against Atf3, Fos, Jun, JunD, Jdp2, and Fra2/Fos2. To achieve AP-1 gain and loss-of-function *in vitro*, mammalian expression plasmids pcDNA3.FLAG-Fos WT (Addgene plasmid #8966) and CMV.Flag-AFOS (Addgene plasmid #33353) was cloned into pENTR.CMV.Flag entry vectors. These entry vectors were then recombined into the pAd/BLOCK-iT™ vector via LR Clonase reactions (Invitrogen) to generate adenoviral vectors expressing CMV-driven cFos (gain-of-function), AP-1 dominant negative A-Fos (loss-of-function). Adenoviral particles were produced by transfecting Pac I-digested pAd plasmids into HEK293A cells using Lipofectamine 2000 (Fisher Scientific 11668027). After overnight incubation, cells were expanded and cultured until approximately 80% cytopathic effect (CPE) developed (10-13 days). Virus particles were collected and amplified. Differentiated C2C12 myotubes were infected with adenovirus, followed by EPS or drug treatment, and then harvested.

### Genetic manipulation of AP-1 in vivo

We used adeno-associated viruses (AAV) and transgenic approaches to control muscle-specific AP-1 expression in mice. For AAV-mediated A-Fos expression, we cloned the A-Fos coding region into the AAV.FLEX vector (Addgene plasmid #28304) and subcutaneously injected into neonatal MLC-Cre mice at 1×10¹¹ viral genomes per mouse. For inducible A-Fos (iA-Fos), we generated dual AAV vectors. The first vector encoded Flag-tagged A-Fos under the TRE-tight promoter (from Addgene #46879), and the second vector expressed rtTAM2 under Cre-dependent control (AAV9.CAG.FLEX.rtTAM2.T2A.GFP) as described^93^. For inducible cFos/JunD overexpression, AAV9.TRE9.JunD.T2A.cFos was constructed and used together with AAV9.FLEX.rtTAM2 vector. To construct the transgenic mouse line for TRE.AP-1 (cFos/JunD), the TRE.JunD.T2A.cFos plasmid was injected into mouse embryos at the one-cell stage through microinjection. PCR-based genotype was used to test the genotype of the resultant pups. For genotyping, genomic DNA was extracted from ear tissue in a buffer containing 670 mM Tris (pH 8.8), 166 mM ammonium sulfate, 65 mM MgCl_2_, 2-mercaptoethanol, and 10% Triton X-100, followed by incubation at 95°C for 10 minutes. Samples were then digested with Proteinase K (10 mg/mL) at 55°C for overnight, inactivated at 95°C for 10 min. TRE-AP1 transgenic mice were genotyped using primers CCGTGCCCTGGCCCACCCTCGTGACC and GAACTCCAGCAGGACCATGTGATCGCGCT. A 509 bp band indicates the presence of the transgene. TRE-JunD.2A.cFos was crossed with Rosa26-LSL-rtTA3 (JAX #029617) and MLC-Cre mice (JAX # 024713) to generate a triple transgenic line allowing DOX-inducible skeletal muscle-specific cFos/JunD expression. The rtTA3 allele was genotyped using mutant primers GCAACGTGCTGGTTATTGTG and AGCTCCAAGAACGAAAAGG, and WT primers AAGGGA GCTGCAGTGGAGTA and CAGGACAACGCCCACACA, following the JAX protocol. Product sizes of 172 and 241 bp were considered as mutant and WT, respectively. The MLC-Cre allele was genotyped using common forward (CACACTGCTCTTCCAAGTGTC), wild-type reverse (AGTTACCTTAATAGCAGACAGATCG), and mutant reverse (GCAAACGGACAGAAGCATTT) primers according to the JAX protocol. The mutant allele yields a 280 bp band; the wild type showed a 200 bp band; and heterozygotes showed both. The PCR cycling (initial denaturation 94°C 5 min; 10 cycles of 94°C 30 s, annealing starting at 65°C and decreasing 0.5°C per cycle for 30 s, extension at 68°C 30 s; followed by 28 cycles at 94°C 30 s, 60°C 30 s, 72°C 30 s; final extension 72°C 5 min) was used for all genotyping.

### Immunofluorescence

Tibialis Anterior (TA) muscles were isolated, embedded in optimal cutting temperature (OCT) compound (VWR 25608-930), and rapidly frozen in dry ice-chilled isopentane. Cryosections of 10-μm thickness were obtained using a Leica (CM1850) cryostat. Sections were permeabilized with 0.3% Triton X-100 in PBS, blocked with 5% normal goat serum, and incubated overnight with primary antibodies against laminin (Sigma L9393), MHC-IIa (2F7; Developmental Studies Hybridoma Bank), and MHC-IIb (BF-F3; Deutsche Sammlung von Mikroorganismen und Zellkulturen) in blocking buffer. Secondary antibodies, including Alexa-647-goat anti-rabbit, Alexa-594-goat anti-mouse IgG, and Alexa-488-goat anti-mouse IgM (Invitrogen, 1:1000 dilution in PBS), were then applied for 1 hour in the dark. Unstained fibers were identified as MHC-IIx based on the composition of mouse TA muscles. Fluorescent images were captured with a Zeiss Axiophot epifluorescence microscope and analyzed using Axiovision software to quantify MHC isoform distribution as a percentage of muscle fiber types, with nuclei counterstained by DAPI. ImageJ was used for analyzing the diameters.

### Transmission electron microscopy

The muscle was immersion-fixed at room temperature with 2.5% glutaraldehyde and 1.25% formaldehyde in 0.1 M cacodylate buffer (CB) at pH 7.2, then placed on ice for 10 min. Crosslinked tissue was then hand-dissected into ∼2mm strips, further fixed overnight at 4 °C, and cut into cubes for fixation in 1.0% OsO4 for 1 hr on ice. Tissue was then washed in CB, followed by an ascending ethanol series to 100% ethanol, followed by propylene oxide, and then infiltrated with epoxy resin (50:50/then 100% resin), hardened in a 60 °C oven overnight, followed by ultramicrotomy, and sections were heavy metal contrasted in lead and uranyl salts before imaging. Images were taken at 50,000x magnification using a transmission electron microscope (JEOL JEM-1400) equipped with a Gatan Orius SC1000 camera. Mitochondrial quantifications were analyzed using ImageJ and Qpath software. Mitochondrial volume density was assessed using the stereological point counting method as desribed^94^, which involves calculating the total area occupied by mitochondria divided by the total area of the field.

### Chromatin immunoprecipitation-sequencing (ChIP-seq)

For chromatin immunoprecipitation (ChIP), approximately 80 mg of quadriceps skeletal muscle tissue was homogenized, cross-linked with formaldehyde, and lysed to extract chromatin. The chromatin was then fragmented to an average size suitable for ChIP-seq analysis (typically around 200-500 base pairs). ChIP was performed using cFos antibodies (Cell Signaling Technology, 2250) in a procedure as described^95^. cFos ChIP DNA and a proportion of total input DNA were used to generate sequencing libraries using the NEBNext Ultra II DNA Library Prep Kit for Illumina (New England Biolabs). Libraries were quantified by Qubit fluorometry, checked for size distribution on an Agilent TapeStation, and sequenced on the NovaSeq 6000 platform to obtain approximately 30 million paired-end 150 bp reads.

### ChIP-seq data analysis

The raw sequencing reads were first evaluated using FastQC (v0.11.9) (https://www.bioinformatics.babraham.ac.uk/projects/fastqc/) to assess overall data quality, including per-base sequence quality, GC content, and adapter contamination. High-quality reads were aligned to the *Mus musculus* reference genome (mm10) using Bowtie2 (v2.4.5) with default parameters to ensure accurate and sensitive mapping^96^. The resulting SAM files were converted to BAM format, sorted, and indexed using samtools (v1.15)^97^. PCR duplicates were removed using samtools rmdup to avoid artificial signal amplification. Peak calling was performed using MACS2 (v2.2.7.1) with a significance threshold set to p-adjusted < 0.05, yielding high-confidence enriched regions in narrowPeak format. ChIPseeker (v.1.28.3), an R package, was used to perform peak annotation^98^. To improve data clarity, ENCODE-defined mm10 blacklist regions were removed from peak files using BEDTools (v2.30.0) to eliminate artifacts and false-positive peaks^99^. Normalization of signal intensities was performed using bam Coverage (v3.5.1) from the deepTools suite, generating coverage tracks in BigWig format. The effective Genome Size parameter was set for mm10 (2150570000) to account for mappable regions during normalization. Enrichment patterns across genomic regions were visualized using heatmaps generated with compute Matrix and plot Heatmap functions in deepTools^100^. The differential peak analysis was performed using DiffBind and the genes/regions were annotated by utilizing ChIPseeker^101,102^. Finally, peaks and signal tracks were visually inspected in IGV (Integrative Genomics Viewer, v2.15.2) to confirm enrichment at loci of interest and to compare peak distributions across samples^103^. The color gradient in IGV represents the frequency or intensity of the ChIP signal, ranging from minimum to maximum signal strength detected across the genome.

### RNA-seq

For RNA-seq, RNA was isolated from mouse muscles using TRI reagent (Sigma T9424-200ML) and reverse-transcribed to cDNA using a cDNA Synthesis kit (170-8891, Bio-Rad). RNA samples were assessed using the Bioanalyzer 2100 system (Agilent Technologies). Libraries were prepared using dUTP in second-strand synthesis, followed by end repair, A-tailing, adapter ligation, USER enzyme digestion, amplification, and purification. Libraries were quantified using Qubit, real-time PCR, and Bioanalyzer. Samples from exercise vs rest groups in WT mice were sequenced using Illumina HiSeq 2000. Samples from the iA-Fos vs control and iOE vs control groups were sequenced on the BGISEQ-500 platform. Base-calling was performed using the ISEQ-500 system. iA-Fos vs control generated approximately 5.26 GB of data per sample with a genome mapping rate of 94.99% and gene mapping rate of 45.95%. In comparison, cFos/JunD overexpressed (iOE) vs control samples generated 6.38 GB per sample with 94.65% genome and 55.08% gene mapping rates. Reads were filtered to remove low-quality, adaptor-polluted, or undetermined sequences. Mapping was performed with HISAT2, and transcript annotation and novel gene discovery were performed using BGI’s internal pipeline. UCSC bigWig and peak files were generated using HOMER. Gene-level read counts were generated using *featureCounts* v1.5.0-p3, and expression was normalized as FPKM. Differential expression analysis was performed using *DESeq2* (v1.20.0), with q-value ≤ 0.05 and |log_2_(fold-change)| > 0.58 as the threshold for significance. Gene ontology (GO) and KEGG enrichment analyses were conducted using DAVID (v6.7). Heatmaps and z-score visualizations were generated using SRPlot and Morpheus.

### GRO-seq

GRO-seq was performed to profile nascent transcription as described previously^104^. Briefly, nuclei were isolated from frozen quadriceps muscle tissues and incubated in nuclear run-on reactions containing Br-UTP for 5 minutes at 30°C to label newly synthesized RNA. After labeling, samples were incubated at 37 °C for 1 hour in the presence of PNK buffer (0.5 M MES, 50 mM MgCl_2_, 50 mM β-mercaptoethanol, 1.5 M NaCl, 10 U SUPERase-In) to stabilize transcripts. Br-UTP incorporated transcripts were enriched using anti-BrdU antibody-conjugated magnetic beads. Beads were pre-blocked and washed before binding, and bound RNA was sequentially washed using buffers of increasing stringency to reduce background. Following enrichment, the RNA was polyadenylated using *E. coli* poly(A) polymerase and reverse transcribed to generate cDNA. Excess primers were removed with exonuclease I, and RNA was degraded by alkaline hydrolysis, followed by neutralization. The resulting single-stranded cDNA was circularized using CircLigase and then re-linearized with APE1 to enable strand-specific library preparation. Libraries were amplified via PCR, and quality was checked prior to sequencing. Run-on reactions from five mice per group were pooled to generate a single library, and all library preparations were carried out in parallel to minimize batch effects.

### GRO-seq data analysis

Nascent RNA from quadriceps muscle of WT and HDAC3-KO exercise vs rest samples was sequenced on the HiSeq 2000 platform with single-end 50 bp reads. Reads were aligned to the Mus musculus reference genome (mm9) using Bowtie2 v2.5.4^96^. BigWig and peak files were generated using HOMER2 for downstream analysis^105^. The identification and quantification of eRNAs were carried out according to previously defined procedures^104^. eRNAs with FPKM > 10 and fold-change > 2 in exercise vs. rest samples were considered differentially expressed. Only bidirectional eRNAs were retained after filtering out sense-strand intragenic eRNAs.

### Kinetic metabolic testing, energy expenditure, and isotope tracing

For glucose tolerance tests (GTT), mice were fasted for six hours with free access to water, and blood glucose levels were measured after an i.p. injection of 2g/kg glucose using the OneTouch Ultra2 glucometer at different time points. To quantify glucose clearance over time, the area under the curve (AUC) was calculated using the trapezoidal rule “AUC = Σ from i = 1 to n–1 [ (Gi + Gi+1) / 2 × (ti+1 – ti) ]” where *Gi* and *Gi+1* are the blood glucose concentrations at time points *ti* and *ti+1*, respectively. Indirect calorimetry was conducted using an automated Comprehensive Lab Animal Monitoring System (Oxymax/CLAMS-HC, Columbia Instruments) to monitor the mice’s metabolism, including energy expenditure, caloric intake, body temperature, and locomotor activity. Briefly, the mice were housed individually and acclimated for 2 days in a metabolic chamber equipped with a running wheel. Quantitative measurements of oxygen consumption (VO2), and carbon dioxide production (VCO2) were recorded every 20 minutes over 72 hours. Food intake and locomotor activity were measured. For glucose uptake analysis, cells or isolated tissues were incubated with 2-[3H(N)] deoxy-D-glucose for 10 minutes, followed by a 6-hour serum starvation period. Cells were then washed three times with cold PBS and lysed in RIPA buffer containing 1% SDS. For tissue samples, muscles were incubated in media containing radiolabeled glucose analog, after which the media was collected to measure radioactivity in disintegrations per minute (dpm) using a scintillation counter (Beckman Coulter LS 6500). Radioactivity values were normalized to protein concentration in cell lysates or tissue samples.

### Muscle performance and contractile physiology

Forelimb maximal muscle grip strength was determined as reported previously^106^. Mice were briefly placed at the base of the grid, allowing them to grip the grid wire and pull backward horizontally until their grip was released. Multiple readings with a 5-minute rest interval were recorded, and the average was calculated. For the wire-hanging test, skeletal muscular strength was measured as described previously^107^. The mice’s forelimb paws were suspended horizontally on a 1 mm-wide, 2 mm-long metallic wire at a sufficient height, with proper bedding. The latency period, the time mice withstand to fall, was recorded, and similar multiple trials per session tests were performed weekly. The average of the trials was regarded as average performance for each session. The treadmill graded exercise test was performed using an Exer 3/6 Treadmill with an electrical stimulus system and the automatic stimulus detection unit (Columbus Instruments) as previously described^107^. Mice were motivated to run by electric shocks at 1 Hz with a stimulation intensity of 3, without inclination. Mice well tolerated the shocks at such high intensity and frequency. No mice showed injury after testing or training. After mice were placed on the stationary treadmill for 5 min, the treadmill started at 6 m/min, the speed was increased by 2 m/min every 10 min, then 30 min for 18 m/min, 15 min for 20, 22, 24, with no further increase in speed beyond 26 m/min. Exhaustion was defined as the time at which the mouse reached 50 cumulative shocks during the entire test. Once the cumulative shocks for a specific lane reached 50, the shocking unit for that lane automatically turned off. At this point, the mouse in that lane could rest on the shocking grid while the treadmill continued running. Mice were then removed from the treadmill after all lanes had finished. For in-vivo muscle physiology, fatigue resistance was determined by measuring muscle dorsiflexor contractile function on TA muscles using the Aurora 1300A 3-in-1 system supported with 610A Dynamic Muscle Control v.5.501 software as previously reported^107^. The hind limb hair of mice was shaved to facilitate efficient muscle location, followed by anesthesia induction using 2.5% isoflurane supplemented with a 1 L/min oxygen flow. The electrodes were positioned on the TA muscle beneath the skin (2-4 mm depth), and the current required to achieve the maximum possible torque was recorded. Maximal tetanic contractions were obtained using a stimulation frequency of 100 Hz for 500 msec. Five-minute rest between two tetanic contractions was allowed to ensure muscle recovery. To induce fatigue, muscles were stimulated with 40 Hz pulses lasting 300 ms every 5 seconds for up to 90 contractions. The fatigue index was expressed by the percentage of maximum force (% maximum).

Ex vivo muscle physiology was performed on isolated EDL muscles^15,108^ using an Aurora Mouse 1200A system with dynamic muscle control v.5.415 software. EDL muscles were continuously immersed in oxygenated Ringer’s solution containing 100 mM NaCl, 4.7 mM KCl, 3.4 mM CaCl2, 1.2 mM KH2PO4, 1.2 mM MgSO4, 25 mM HEPES, and 5.5 mM d-glucose, and maintained at 24 °C throughout the experiment. A single stimulus with a duration of 0.2 ms was applied for twitch stimulation and repeated at a frequency of 120 Hz for 500 ms for measuring tetanic maximal force generation, followed by 5 min recovery between two tetanic contractions. For fatigue induction, muscles were stimulated every second for 8 min using 40-Hz pulses lasting 330 ms. The fatigue index was expressed as the percentage difference in force between the first contraction and each subsequent contraction for EDL, or as the percentage of force remaining after the preceding contraction. A burst of 50 maximal tetanic contractions (120 Hz for 500 ms) was applied to ensure complete fatigue across all samples. The recovery protocol started 1 s after the last burst contraction. A maximal tetanic stimulation (120 Hz for 500 ms) was given every 5 min for 30 min, and the force recovery was expressed as a percentage of the maximal isometric tetanic force.

### Western blot

Samples were lysed in RIPA buffer (50 mM Tris pH 8.0, 150 mM NaCl, 0.1% SDS, 0.5% Sodium deoxycholate, 1% Triton X-100) containing 1% protease inhibitor (Fisher Scientific A32965). Protein concentration was determined by the BCA assay reagent as described in the manufacturer’s (Thermo Scientific-23227). Equal amounts of protein were resolved by SDS-PAGE and transferred onto PVDF membranes (Bio-Rad, 1704273). The membrane was incubated in 5% milk-blocking solution for 2 h at room temperature (RT) and probed against the primary antibody against cFos (Cell Signaling Technology, 2250S), FLAG (Sigma, F4042), β-Actin (Santa Cruz Biotechnology, SC-47778), GFP (CST, 2956T), GAPDH (Santa Cruz, SC-47724), HSP90 (CST, 4874S), ATF3 (VWR, 89294-974), and OXPHOS (catalog number pending), each diluted 1:2000 in TBST. After washing with TBST, the membrane was incubated with HRP-conjugated IgG secondary antibody for 2 h, washed again with TBST, and immunoreactive bands were visualized by chemiluminescence (Lumiquant AC600, ACURONBIO Technology Inc).

### Statistical analysis

All results are expressed as mean ± SEM. Statistical significance was assessed by Student’s *t-test* (unpaired two-tailed) or ANOVA followed by post-Bonferroni’s multiple comparisons whenever required and analyzed using GraphPad Prism (version 8,10; GraphPad Software). A *p-value* of less than 0.05 was considered statistically significant.

## DATA ACCESS

RNA-seq, GRO-seq, and ChIP-seq data are available upon request.

## ACKNOWLEDGEMENT

We thank Dr. Abhinav Jain at MD Anderson Epigenomics Profiling Core for cFos ChIP-seq analysis, Dr. Michael A Mancini and Debra Townley at the Baylor College of Medicine (BCM) Integrated Microscopy Core for electron microscopy analysis, Dr. Kazuhiro Oka, Dr. Corinne Sonnet, and Austin Seal at BCM Gene Vector Core for producing recombinant AAV vectors, Drs. Yefei Wen and Baomei Qian for immunostaining and mouse colony maintenance, and Dr. Pradip Saha at the BCM Mouse Metabolism and Phenotyping Core (supported by UM1HG006348, R01HL130249, and R01DK114356) for CLAMS analysis. UKS was supported by Frank Belton Kimmel and Sandra Kimmel Endowed Post-Doctoral Fellowship and a training fellowship from the Gulf Coast Consortia (GCC) on the Training in Precision Environmental Health Sciences Program (T32ES027801). LY was supported by the National Key Research and Development Program of China (2022YFC2703500) and National Natural Science Foundation of China grants (82371617, 82571836). The study was partially supported by the NIH grant DK111436. The authors are also thankful to the grants AG069966, ES034758, John S. Dunn Foundation, the Texas Medical Center Digestive Diseases Center (P30DK056338), and the Gulf Coast Center for Precision Environmental Health (GC-CPEH, P30ES030285).

## AUTHOR CONTRIBUTION

AC conducted in vivo muscle physiology, gene expression, and biochemistry analyses. AC, TY, and HY conducted mouse experiments, including DOX treatment, in vivo muscle performance, GTT, qPCR, histology, western blot, and electron microscopy. UKS performed ChIP-seq and RNA-seq data analysis. SH performed AP-1 knockdown and overexpression *in vitro* and constructed plasmids. JJ performed GRO-seq. MD and BF performed GRO-seq analysis. CS performed the continuous AP-1 western blot. HT measured ROS. ESO helped with DOX treatment. JP and EB performed ex vivo muscle physiology analysis. LL and JX constructed the AP-1 transgenic mouse line. YL provided financial support for HY. AC, TY, SH, ZH, YL, and ZS interpreted the data. ZS conceptualized, designed, and supervised the project. AC and ZS wrote the manuscript with the input of other authors.

---

**Figure S1.**
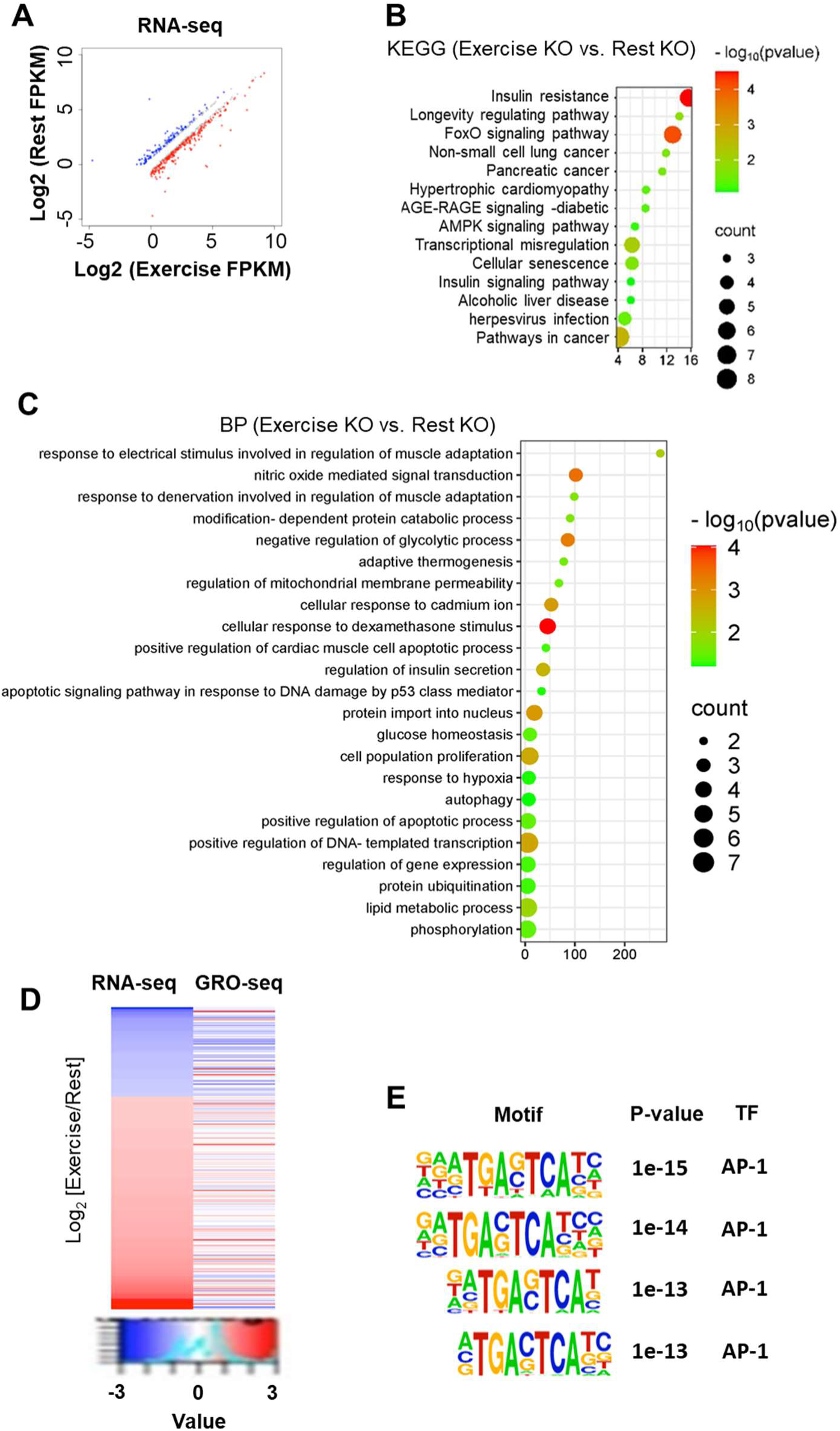
GRO-seq and RNA-seq in muscle-specific HDAC3 knockout mice after a bout of treadmill exercise. **(A)** Scatter plot showing differentially expressed genes (DEGs) in quadriceps from HDAC3 muscle-specific knockout (KO) mice subjected to acute treadmill running compared to resting controls (exercise vs. rest; q < 0.05, |log_2_(fold-change)|>0.58). **(B-C)** Gene Ontology (GO) biological process (BP) and KEGG pathway enrichment of upregulated DEGs (exercise vs. rest) in HDAC3-MKO mice using DAVID analysis and SRplot. **(D)** Heat map of RNA-seq and GRO-seq signals at gene bodies in quadriceps from HDAC3 MKO mice. Values shown as row-wise z-scores calculated from FPKM values. **(E)** Top transcription factor motifs in exercise-induced eRNAs from quadriceps of KO mice.

**Figure S2.**
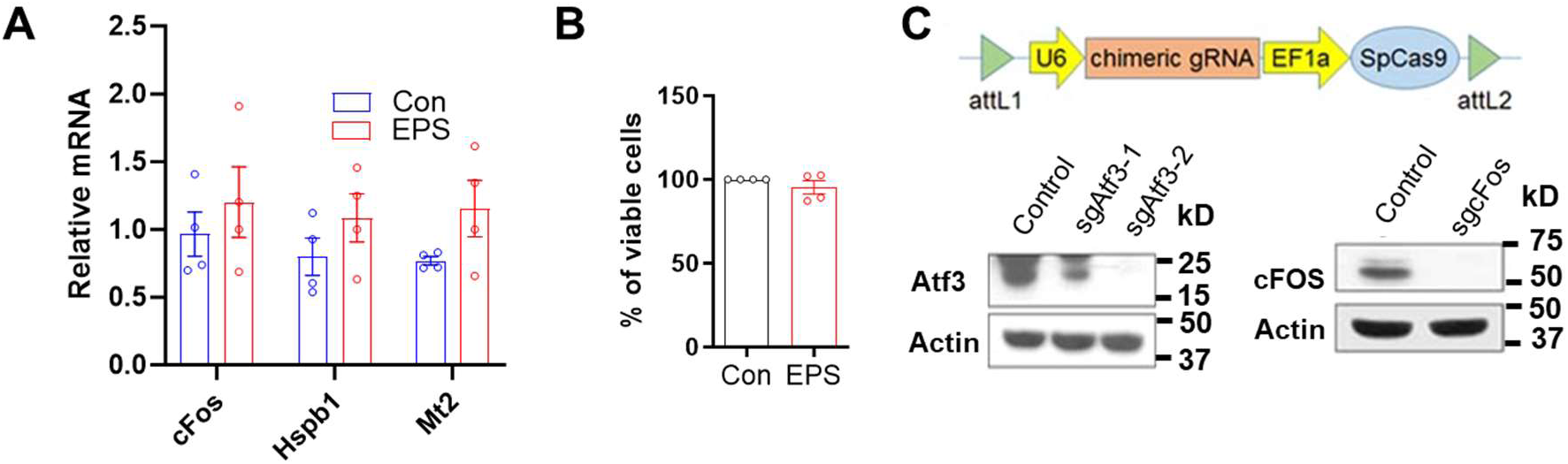
Effects of EPS and AP-1 knockdown in C2C12 myotubes. **(A)** RT-qPCR analysis of undifferentiated C2C12 myotubes after contractions induced by electric pulse stimulation (EPS). n=3-4. (**B)** Cell viability assay to determine the toxicity of EPS in differentiated C2C12 cells, n=3-4. **(C)** Immunoblot analysis of C2C12 myotubes infected with adenovirus expressing sgRNAs for knockdown of individual AP-1 members (Atf3 and cFos).

**Figure S3.**
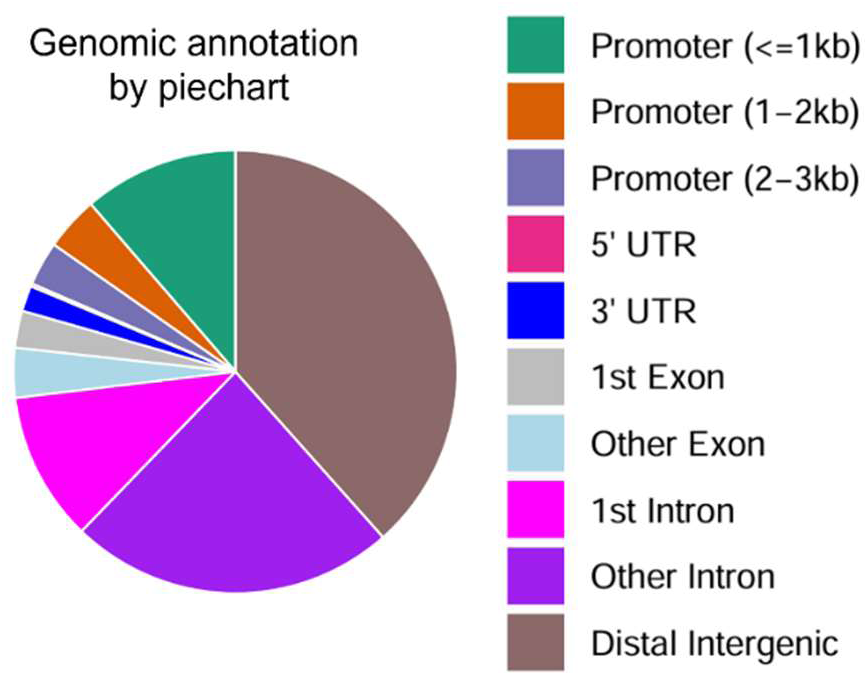
Genomic distribution of cFos ChIP-seq peaks in muscles. Distribution of the upregulated ChIP-seq (963 peaks) in Exercise vs WT muscles.

**Figure S4.**
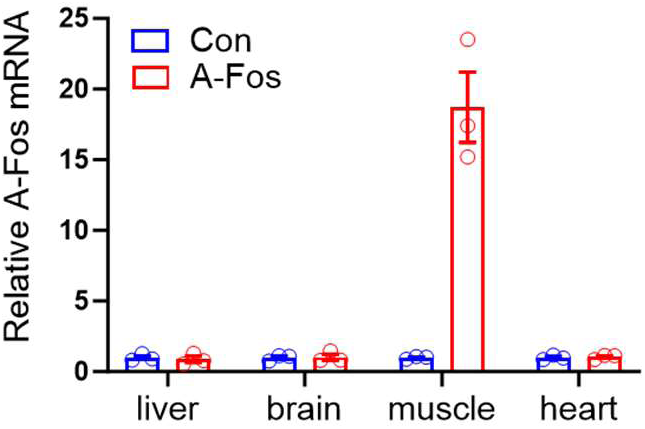
Expression of AAV-mediated expression in muscles. RT-qPCR analysis of different tissues from mice injected with AAV expressing A-Fos. Data are mean ± SEM.

**Figure S5.**
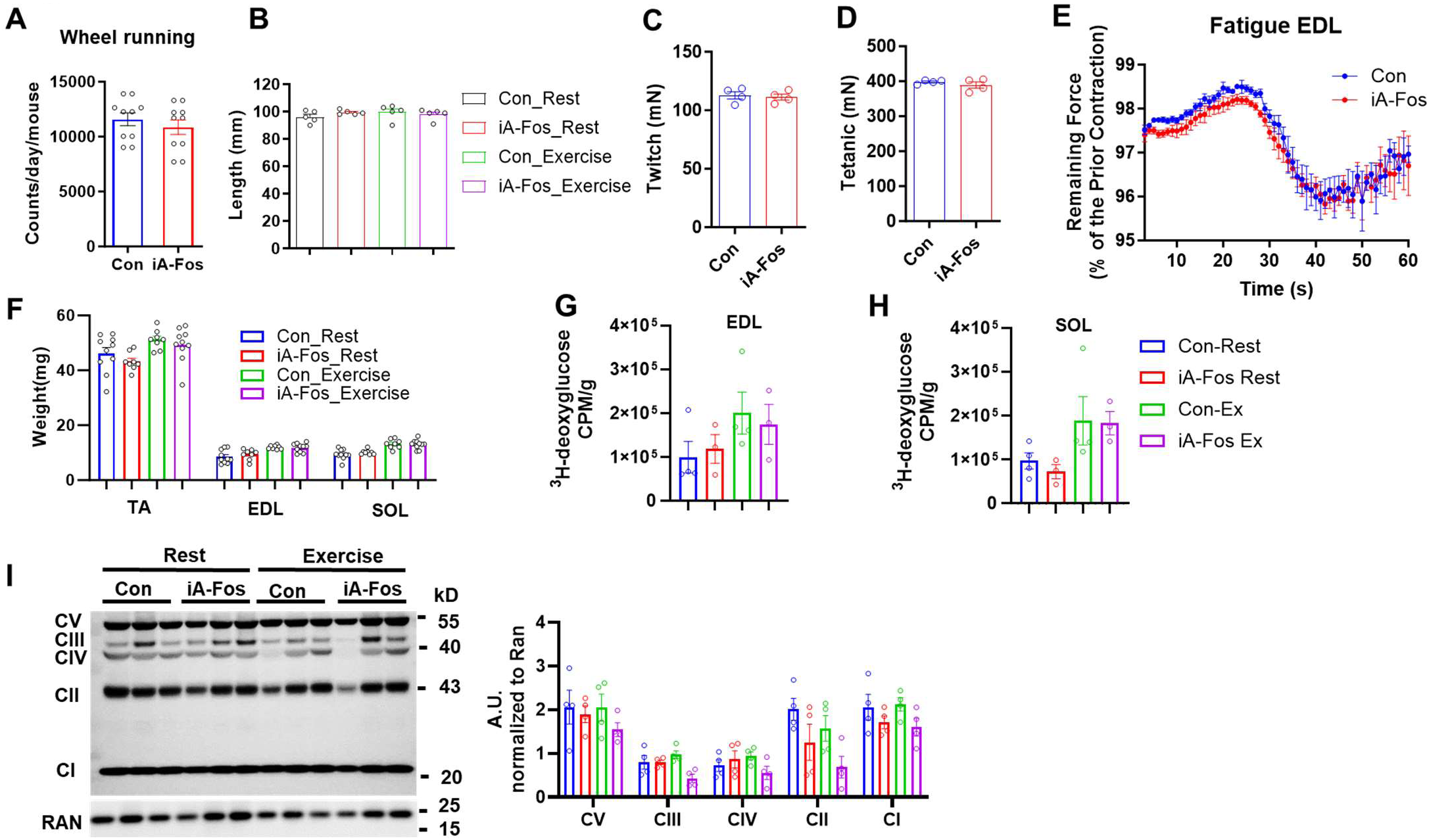
Lack of change in baseline muscle strength, fatigue, or glucose uptake in iA-Fos mice. **(A)** Wheel-running activity in adult mice. **(B)** Body length. **(C-E)** E*x vivo* muscle physiology analysis of twitch or tetanic force and fatigue resistance in the EDL muscles. **(F)** Muscle mass. **(G-H)** The glucose uptake assay with ^3^H-deoxyglucose in isolated EDL and soleus muscles. **(I)** Immunoblot and quantification of oxidative phosphorylation (OXPHOS) protein complexes (complex I-V, CI-CV). Data are mean ± SEM.

**Figure S6.**
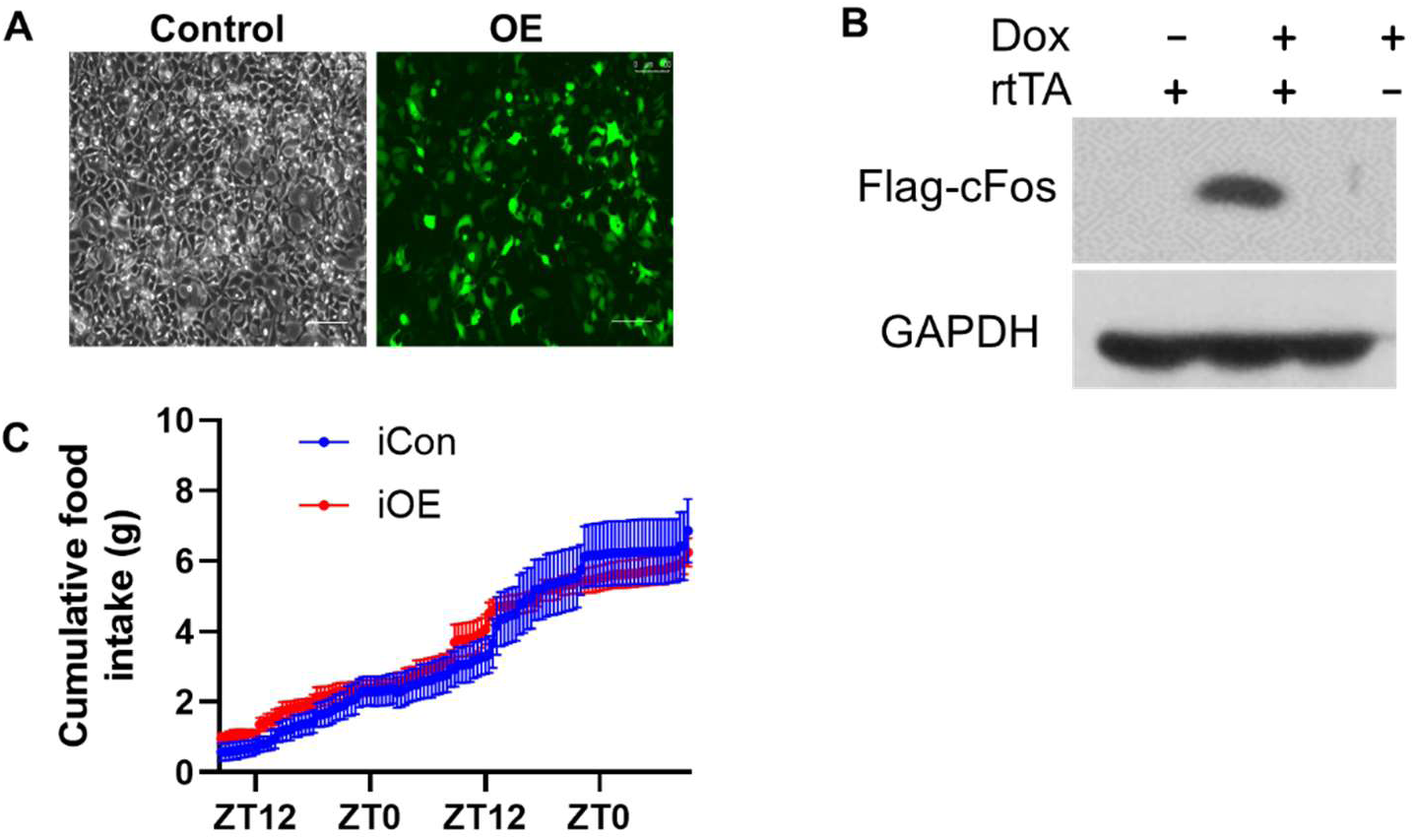
Lack of change in cumulative food intake in iOE mice. Data are mean ± SEM.

**Figure S7.**
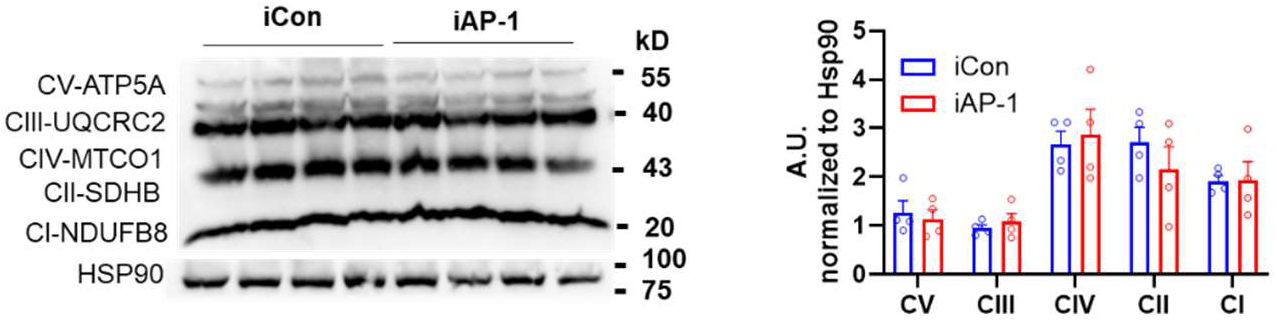
Lack of change in OXPHOS protein levels in iAP-1 mice. Immunoblot analysis of TA muscles.

**Figure S8.**
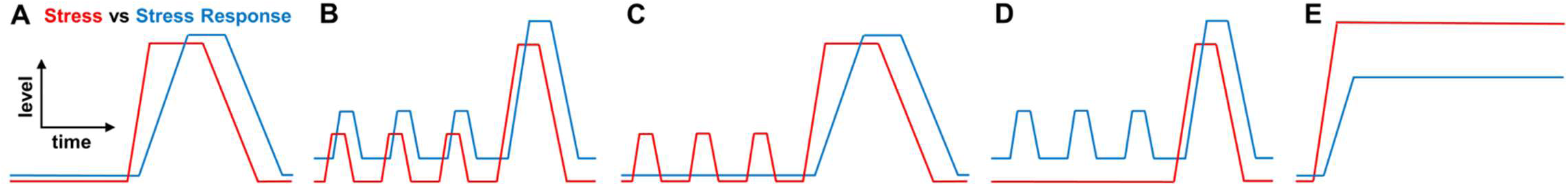
The intricate relationship between stress and stress response in resilience.

**Table S1. DEGs (exercise vs. rest) in WT muscles**

**Table S2. sgRNA sequences for knockout of AP-1 family members**

**Table S3. DEGs (iA-Fos vs. WT control) after acute treadmill exercise**

**Table S4. DEGs (iOE vs. iCon) in mouse TA muscles**

**Table S5. DEGs (cAP-1 vs. cCon) in mouse TA muscles**

